# *Escherichia coli* adapts metabolically to 6- and 7-fluoroindole, enabling proteome-wide fluorotryptophan substitution

**DOI:** 10.1101/2023.09.25.559291

**Authors:** Christin Treiber-Kleinke, Allison Berger, Lorenz Adrian, Nediljko Budisa, Beate Koksch

## Abstract

Nature has scarcely evolved a biochemistry around fluorine. However, modern science proved fluorinated organic molecules to be suitable building blocks for biopolymers, from peptides and proteins up to entire organisms. Here, we conducted adaptive laboratory evolution (ALE) experiments to introduce fluorine into living microorganisms. By cultivating *Escherichia coli* with fluorinated indole analogues, we successfully evolved microbial cells capable of utilizing either 6-fluoroindole or 7-fluoroindole for growth. Our improved ALE protocols enabled us to overcome previous challenges and achieve consistent and complete adaptation of microbial populations to these unnatural molecules. In the ALE experiments, we supplied fluoroindoles to auxotrophic *E. coli* bacteria, exerting strong selective pressure that led to microbial adaptation and growth on monofluorinated indoles. Within the cells, these indoles were converted into corresponding amino acids (6- and 7-fluorotryptophan) and incorporated into the proteome at tryptophan sites. This study is a first step and establishes a strong foundation for further exploration of the mechanisms underlying fluorine-based life and how a formerly stressor (fluorinated indole) becomes a vital nutrient.

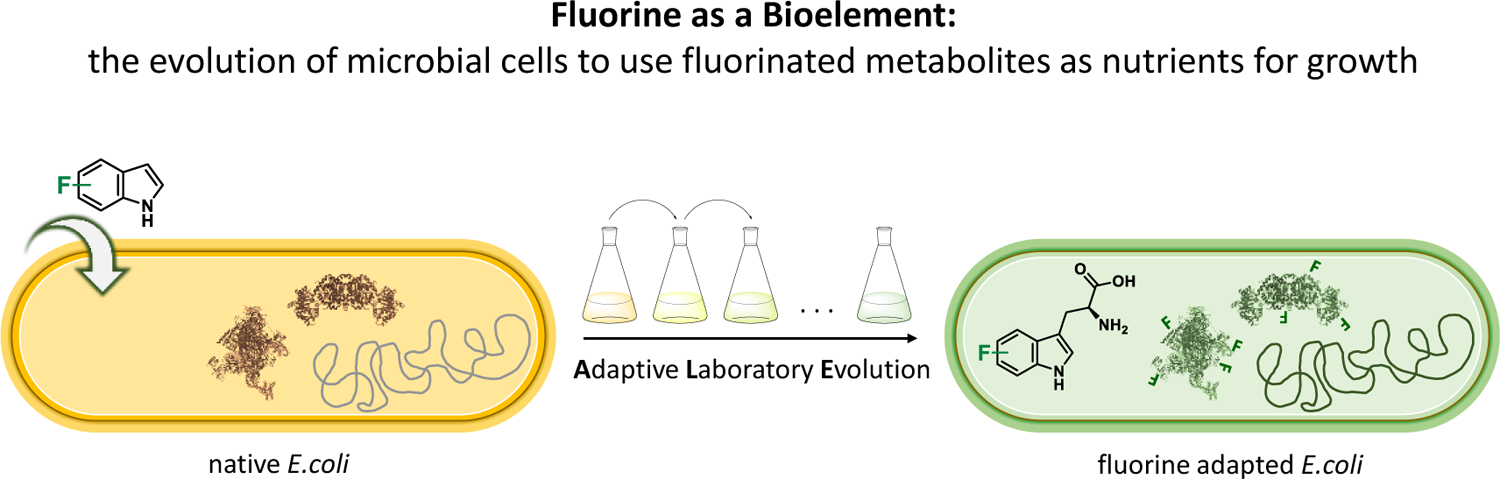

## Introduction

Although fluorine is abundant in the earth’s crust, it is scarcely found in biomolecules. Only a few plants and microorganisms can produce organofluorine compounds.[1,2] One prominent example is the well-studied *Streptomyces cattleya* that biosynthesizes the amino acid 4-fluorothreonine (4FT) and at the same time produces toxic fluoroacetate (FA).[3] In terms of chemistry, this is a challenging task as a C-H bond has to be replaced with a polar-reversed C-F bond. In fact, there is such an enzyme known as fluorinase, that enables C-F bond formation in biological systems.[4,5] The biotechnological potential of such biotransformations is enormous, and consequently, there is much current research in this direction.[6–9]

The paradox of fluorine’s high abundance in an inorganic context yet low abundance in the biomolecular context is discussed on the literature.[10] Fluorine’s inaccessibility for biological systems is attributed to its unique physical and chemical properties, such as the high heat of hydration (∼120 kcal mol^−1^) of the fluoride ion resulting in easy solvation and extreme weak nucleophilicity as well as the very high redox potential (E^0^ = 2.866 V) hampering its oxidation and precludes fluorine from haloperoxidase catalyzed halogenation reactions.[1,11] These are the main reasons discussed why organisms have scarcely been presented with the opportunity to adapt to this extraordinary element. In contrast, the relative natural abundance of other biogenic organohalides, especially organochlorides, is notably higher (several thousand compounds are known) because they were sufficiently present in the prebiotic environment and therefore microbes could evolve to metabolize them (e.g., enzymatic halogenation).[12–14]

Microbes, in particular bacteria, have an astonishing capacity to overcome environmental stress through adaptation, as is observed by the emergence of antibiotic or xenobiotic resistance and adaptation to extreme environments, e.g., extremophilic microorganisms.[15,16] For example, adaptation to substances of anthropogenic origin in polluted environments impressively demonstrates that evolution can take place over relatively short time scales.[17,18] One further aspect of high current relevance is microbe-mediated catabolism of organofluorine compounds, in particular poly- and perfluoroalkyl substances (PFASs) that represent an extremely serious environmental problem due to their persistence.[19]

The natural process of evolution can be realized under controlled laboratory conditions by means of so-called Adaptive Laboratory Evolution (ALE) in which a particular experimental selection pressure is applied to microbes. If the evolved organisms are able to use xenobiotic compounds as nutrients, in the best case incorporate them into biomolecules that can be isolated and studied, this is beneficial from a biotechnology perspective. This explains the growing interest in the community in nature-inspired evolutionary experiments, often with great practical benefits.[20–24] Tryptophan is an attractive target for proteome-wide replacement for three main reasons: it is uniquely encoded by the UGG codon; it has low abundance of about 1 % in the proteome thus only a limited number of proteins are expected to be affected; and the indole moiety provides a great toolbox for chemical interactions.[25] The first experiment of this type was performed with auxotrophic *Bacillus subtilis* cultures grown in minimal media, where tryptophan (Trp) would be replaced by 4-fluorotryptophan (4FTrp) during serial dilution while using an external mutagen to accelerate the adaptation process.[26] This pioneering study was recently repeated with optimized cultivation conditions and with the inclusion of other analogues such as 5-fluorotryptophan (5FTrp) and 6-fluorotryptophan (6FTrp) in the serial dilution ALE experiment.[27,28] In 2001, Bacher and Ellington revisited earlier Wong experiments with 4FTrp-tolerant *Escherichia coli* strains[29] and provided the first genomic studies; unfortunately minute amounts of natural Trp (about 0.03 %, 90 nM) were present in fully adapted cultures.

The central problem of these prior studies is their reliance on the chemical synthesis of the fluorinated tryptophan analogues, which always leaves traces of natural Trp which narrows the selective pressure and might subvert the adaptation process. Thus, indole and its derivatives as small precursors can be advantageous because they can be obtained commercially at extremely high degrees of purity. The suitability of this method was first demonstrated by Hoesl *et. al.* and more recently by our lab. [30,31] Our 2021 study on the adaptation of *Escherichia coli* to 4-fluoroindole (4Fi) and 5-fluoroindole (5Fi) showed via multi-omics analysis that the two evolved populations were remarkably distinct with respect to the observed rearrangements in regulatory networks, membrane integrity, and quality control of protein folding. [31]

This finding motivated us to expand our ALE experiments to include 6-fluoroindole (6Fi) and 7-fluoroindole (7Fi), which are isosteric but have distinctly different physicochemical and photophysical properties.[32,33] We have further improved the overall experimental setup for the ALE procedure and provided many important parameters that are of general importance in attempts to get bacteria fully “addicted”[34] to amino acid analogues and noncanonical amino acids in general. This is confirmed by our extensive ALE studies on Trp-auxotrophic *E. coli* stains, which produced highly adapted bacterial lines capable of growing exclusively on these xenobiotics, accompanied by a clean, proteome-wide substitution of Trp by either 6FTrp or 7FTrp.

## Results

### Metabolic configuration of cells

Stringent *Escherichia coli* auxotrophism towards Trp is a prerequisite for successful ALE experiments with fluoroindole. Therefore, the endogenous pathways of Trp synthesis must be knocked out in the genome. In this particular case, Trp auxotrophy was produced by deleting genes of Trp biosynthesis (*trpLEDC*; trp operon) and the Trp degradation pathway (*tnaA*; tryptophanase enzyme), while the genes for tryptophan synthase (TrpS; *trpBA*) and tryptophanyl-tRNA synthetase (TrpRS; *trpS*) remained in the genome (Figure S4). The metabolic prototype strain *E. coli* MG1655 (Δ*trpLEDC* Δ*tnaA*) was designated as TUB00 (**Figure 1**). The underlying principle of metabolic processing of F-indole and incorporation of FTrp is based on the substrate promiscuity of the two enzymes TrpS[35–37] and TrpRS[26,29,38] (Figure S5). Since the metabolic sources of indole (*tnaA*) and Trp (*trpLEDC*) are eliminated the ALE experiment requires bacteria to accept indole or a fluorinated indole analogue and produce the corresponding amino acid via TrpS and L-serine to survive. Thus, depriving the cells of the canonical substrate (indole) while supplying them continually with the fluorinated indole analogue exerts selective pressure and brings about the adaptation process.

**Figure 1.**
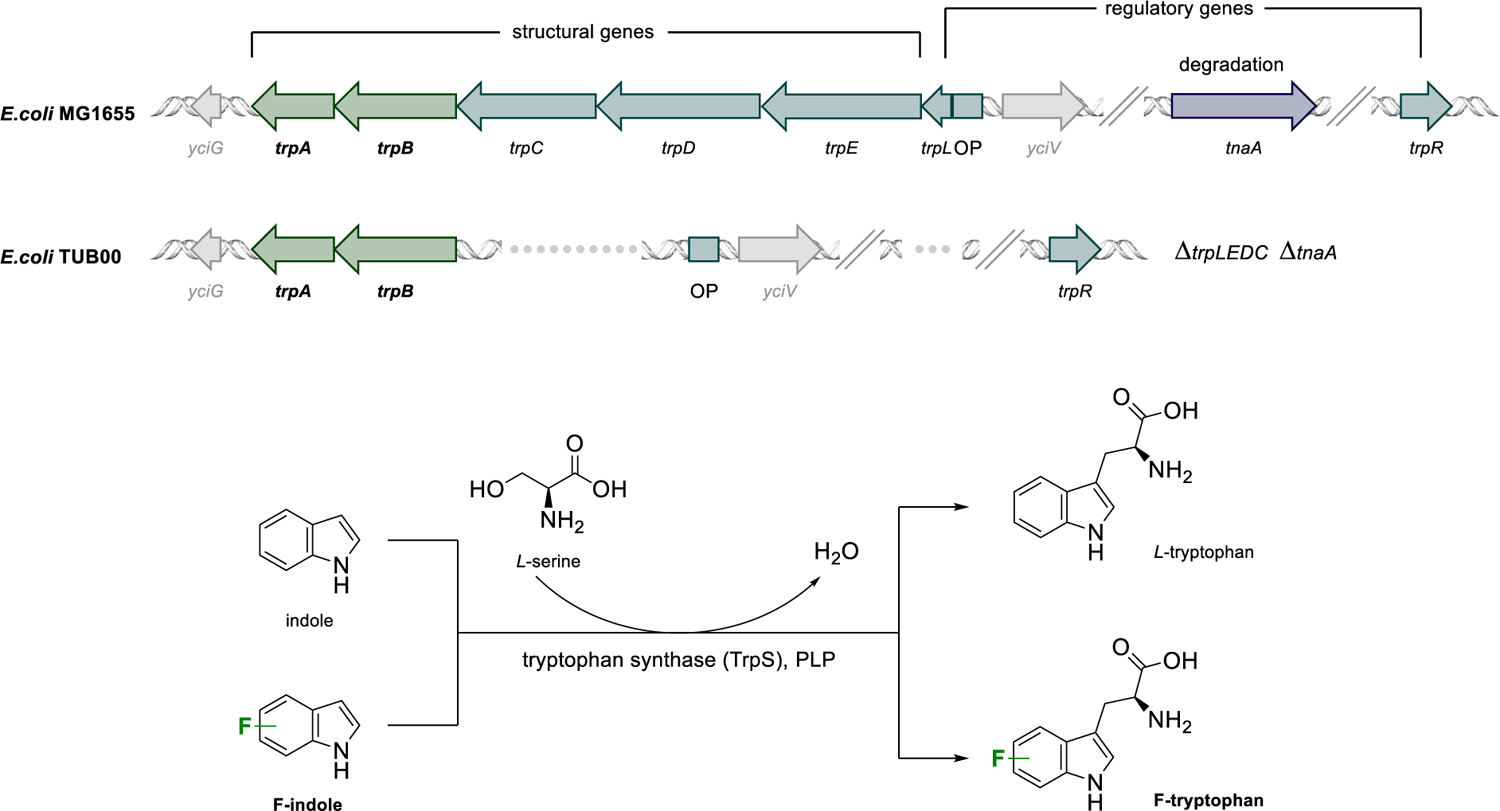
Upper panel: Genes encoding Trp biosynthesis and degradation in *E. coli* parental strain MG1655 and Trp-auxotrophic strain TUB00 (Δ*trpLEDC* Δ*tnaA*).[39] Lower panel: Tryptophan synthase (TrpS) encoded by *trpBA* uses pyridoxal-5’-phosphate (PLP) as cofactor and L-serine as substrate to catalyze the intracellular production of Trp from indole or FTrp from F-indole.

### Design of adaptation experiments

A conceptual overview of the ALE experiments we performed is shown in Figure 2. Four parallel experiments were designed: 1) a positive “indole control”, 2) a positive “Trp control”, 3) adaptation to 6Fi and 4) adaptation to 7Fi. The controls serve to determine fluorine-induced effects and general consequences of long-term cultivation, respectively. We opted to conduct the experiments by allowing liquid cultures to grow under a given set of conditions until they reached early stationary phase; once this occurred, a constant cell number corresponding to an OD_600_ value of 0.02 was used to inoculate the subsequent culture (defined as a “passage”). Unlike the alternative approach of “continuous cultivation”, this kind of serial transfer regime is characterized by alternating “feeding-and-starvation” regimes or “seasonal environments”[40] (i.e., a population bottleneck), but benefits from its flexibility when adjustments to cultivation conditions are needed.[41] Furthermore, we chose the serial transfer approach because this type of experimental management is closest to natural selection in that it mimics changing conditions in the real environment of natural populations. Consist inoculation of each new liquid culture with a normalized amount of cells ensures that even poorly growing cultures are not suffer an evolutionary disadvantage (e.g., our primary goal was not to select for rapidly growing populations).[42]

**Figure 2.**
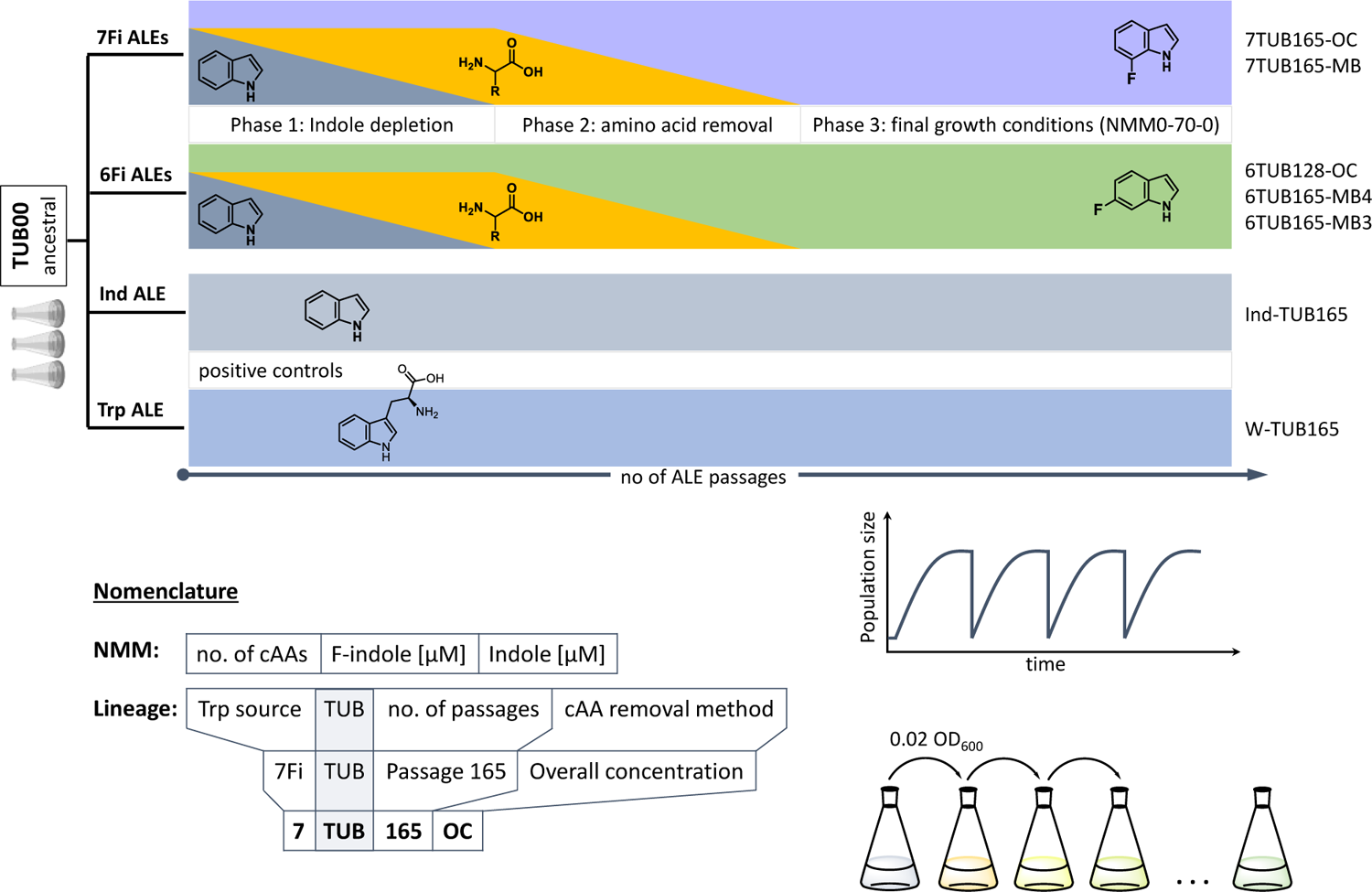
Conceptual ALE trajectories. Four parallel evolution experiments on populations/lineages over the course of a number of passages under different growth conditions were conducted: two positive controls (indole and Trp, lower grey bars) and adaptation to 6Fi (green middle cartoon) or 7Fi (purple upper cartoon). Molecular structures of indole, Trp, 6Fi and 7Fi are shown for the respective experiment. Increasing passage number (independent variable) is indicated by lowest black right arrow; lineage nomenclature follows Z-TUB-X-OC/MB pattern where “Z” indicates the Trp-source (Ind, Trp, 6Fi or 7Fi), “X” the passage number and “OC/MB” the cAA removal method; growth condition nomenclature follows NMM(a-b-c) pattern where “a” indicates the number of cAAs, “b” the concentration of F-indole [µM] and “c” the concentration of indole [µM]. Phases 1 (indole depletion), 2 (amino acid depletion) and 3 (adaptation in presence of constant concentration of only 6Fi or 7Fi as supplement) are indicated. Phase 2 realized according to overall concentration (OC) or metabolic block (MB) strategies (see text for details).

Three biological replicates of the starting point strain TUB00, which we assumed to be isogenic, were allowed to proliferate in synthetic minimal medium (New Minimal Medium, NMM) containing essential nutrients such as phosphate, ammonium, glucose, and vitamins. NMM was differentially supplemented as follows. Indole control populations grew in NMM with a constant concentration of 70 µM indole and high concentration of 19 of the 20 canonical amino acids (cAA); that is, all expect Trp. Trp control populations grew in NMM with no indole but a constant concentration of 70 µM Trp and high concentration of the 19 cAAs (cultivation schemes shown in Figure S7). Populations adapting to 6Fi and 7Fi grew in NMM variably enriched (see below) with indole, the 19 cAAs, and the respective fluorinated indole (see Tables S1-S4 for medium compositions). Growth conditions were abbreviated in all cases as NMM([**x**]**a**-**b**-**c**), where **x** is the concentration of cAAs present in mg/L (only indicated when the concentration was below 50 mg/L); **a** is the number of different cAAs present; **b** is the concentration of 6Fi or 7Fi in µM; and **c** is the concentration of indole in µM (Figure 2). To ensure the purity of the purchased 6Fi and 7Fi precursors, we determined the level of parent indole in batches by GC-MS and, indeed, indole was not detected within the limits of the method (see Figures S1-S3).

The three phases of adaptation to 6Fi and 7Fi are also represented in Figure 2. In phase 1, indole concentration is gradually, stepwise reduced to zero. In phase 2 the 19 cAAs are gradually, stepwise removed according to two distinct strategies (see paragraph below) from the growth medium to exclude possible contamination by traces of indole or tryptophan, which would disturb the adaptation process.[29] Phase 3 represents the period of evolution in which only the fluorinated indole analogue is present as NMM supplement. As noted above, the adaptation experiments designed here proceeded according to “passages” and it is important to note that the criteria that a population had to meet in order to be subjected to the subsequent growth condition were as follows: the population had to have reached early stationary phase under the given condition and proceeded through at least two full passages under that given condition; in some cases more than 2 passages were required to observe stable growth, or no further growth improvement was observed. This is why the number of days of growth does not correspond 1:1 with passage number. Only after this criterion was met were the conditions made more stringent to increase the evolutionary pressure being placed on the population.

Phase 2 was designed according to two different approaches: so-called “overall concentration (OC)” or “metabolic blocks (MB)”. OC and MB are visually represented in Figure 3. In the OC approach, the overall concentration of all 19 amino acids was equally reduced gradually and stepwise from 50 to 0 mg/L. The removal of amino acids in the MB approach was not based on their concentration, but rather on their origin in core metabolism (i.e., their metabolic identity). Canonical amino acids were also assigned to metabolic blocks considering factors such as their metabolic cost in terms of energy and precursor requirements.[43,44] Whereas we and others successfully applied the MB approach in previous adaption experiments,[30,31] the OC approach presented here represents an innovation. We thought that the gradual reduction of all amino acids is more gentle to cells than the sudden elimination of whole metabolic blocks. In this way, cells are softly encouraged to turn on their amino acid biosynthesis processes instead of being abruptly confronted with ribosome stalling and mistranslation effects likely caused by sudden amino acid starvation.[45] Lineage names were assigned according to the pattern Ind-TUBX, Trp-TUBX, 6TUBX-OC, 6TUBX-MB4, 6TUBX-MB3, 7TUBX-OC, or 7TUBX-MB where “Ind” stands for positive indole control, “Trp” stands for positive Trp control, “6/7TUB” stands for the respective fluoroindole adaptation, “X” is the passage number, and “OC” and “MB” stand for the cAA removal strategy used.

**Figure 3.**
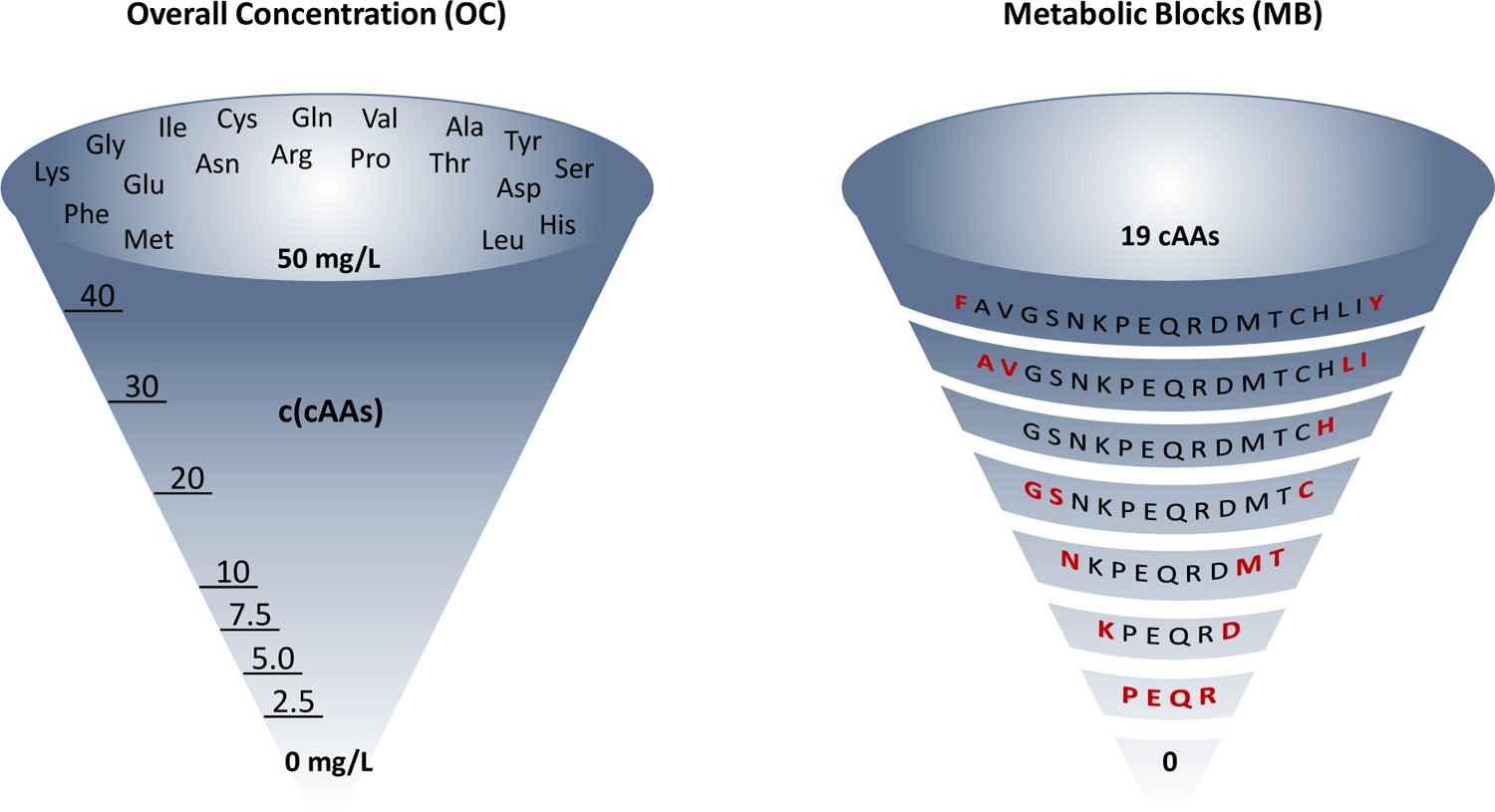
Graphical representation of the differences between OC and MB approaches for amino acid removal in ALE Phase 2, which is a general design concept of our ALE experiments. OC: mixture of 19 amino acids stepwise reduced in concentration from 50 mg/L to 0 mg/L. MB: 19 amino acids removed by identity (shown in red), as metabolic blocks or one by one (in case of the 7Fi ALE). This is a representative example (6TUB128-OC adaptation for the OC approach and 6TUB165-MB4 adaptation for the MB approach) but does not reflect the experimental details for all ALE experiments (shown below).

### Adaptive laboratory evolution experimental setup for 6-fluoroindole (6Fi)

Overall, Trp-auxotrophic *E. coli* tolerated 6Fi remarkably well (Figures 4-6). In phase 1, cells responded sensitively to the challenge of indole depletion, as evidenced by strongly fluctuating OD_600_ values. However, after the populations ceased to be dependent on indole, cell growth remained stable and at a high level.

**Figure 4.**
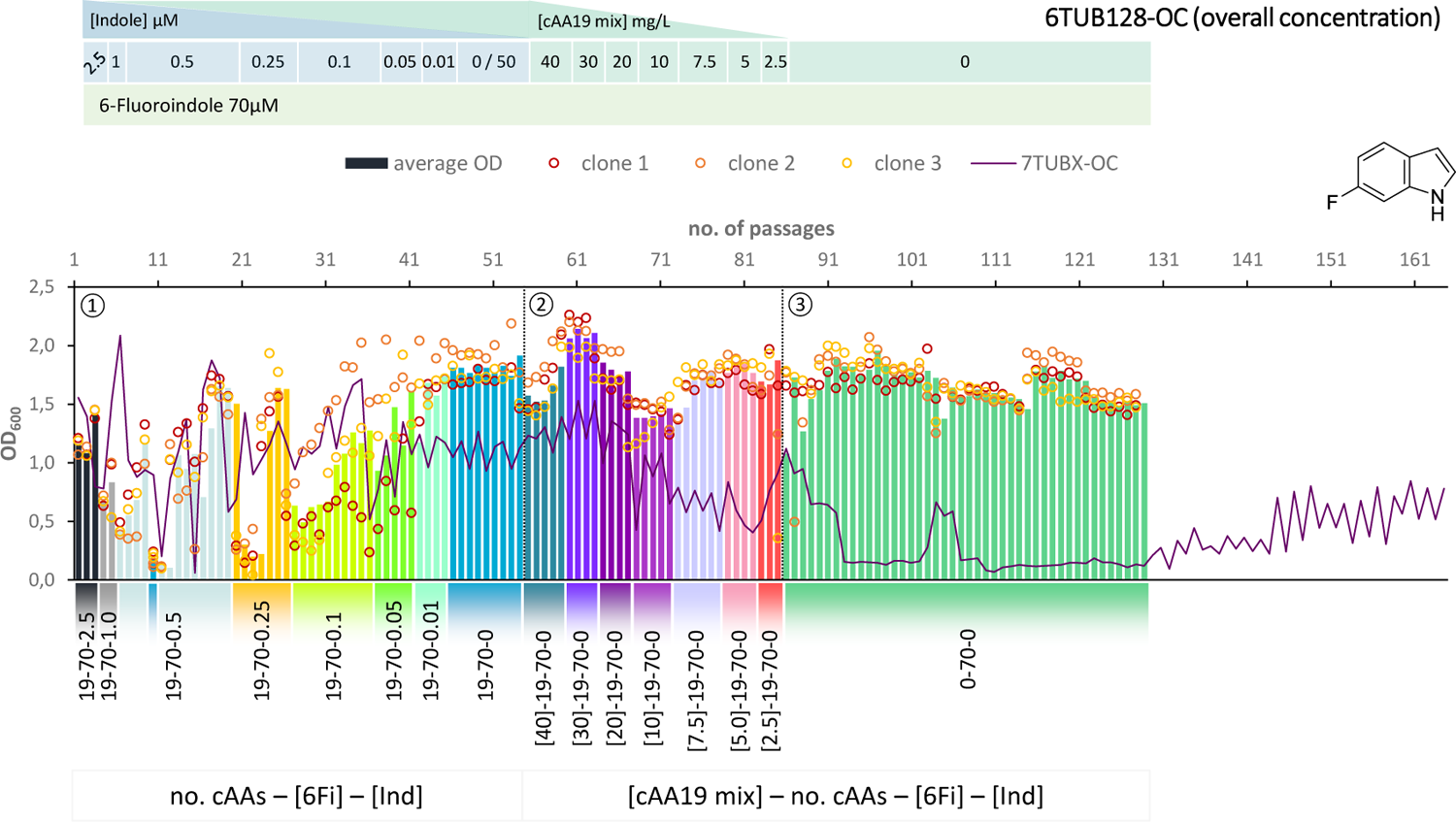
Cultivation scheme of the 6TUB128-OC lineage. Optical density (OD600) is plotted against the number of passages (reinoculation steps); average OD600 is shown as bars and individual clones are shown as circles. The corresponding 7TUBX-OC counterpart is depicted as purple line. ALE phases ①, ②, ③ and medium composition are illustrated; in the OC approach the concentration (mg/L) of the cAA19 mix is indicated in squared brackets [cAA conc.]-a-b-c.

Surprisingly, the OC and MB approaches give approximately equal results in terms of adaptation speed and success. Adaptation took place in a similar time frame (53 passages in the mutual phase 1 and in phase 2 for OC: 31 passages (Figure 4), for MB: 28 passages) and the different lineages show equally good growth behavior. When the amino acid supply in the MB-lineage was almost depleted (9 cAAs left, at passage 73) we tried the removal of different metabolic blocks, which yielded separate lineages 6TUB165-MB4 (Figure 5) and 6TUB165-MB3 (Figure 6). Although the cultivation schemes of the MB-lineages show large variations in cell densities upon removing the amino acid blocks “AVLI” and “NMT” (MB4) as well as “PEQR” (MB3), even removing different blocks was useful here, and the cells recovered very quickly.

**Figure 5.**
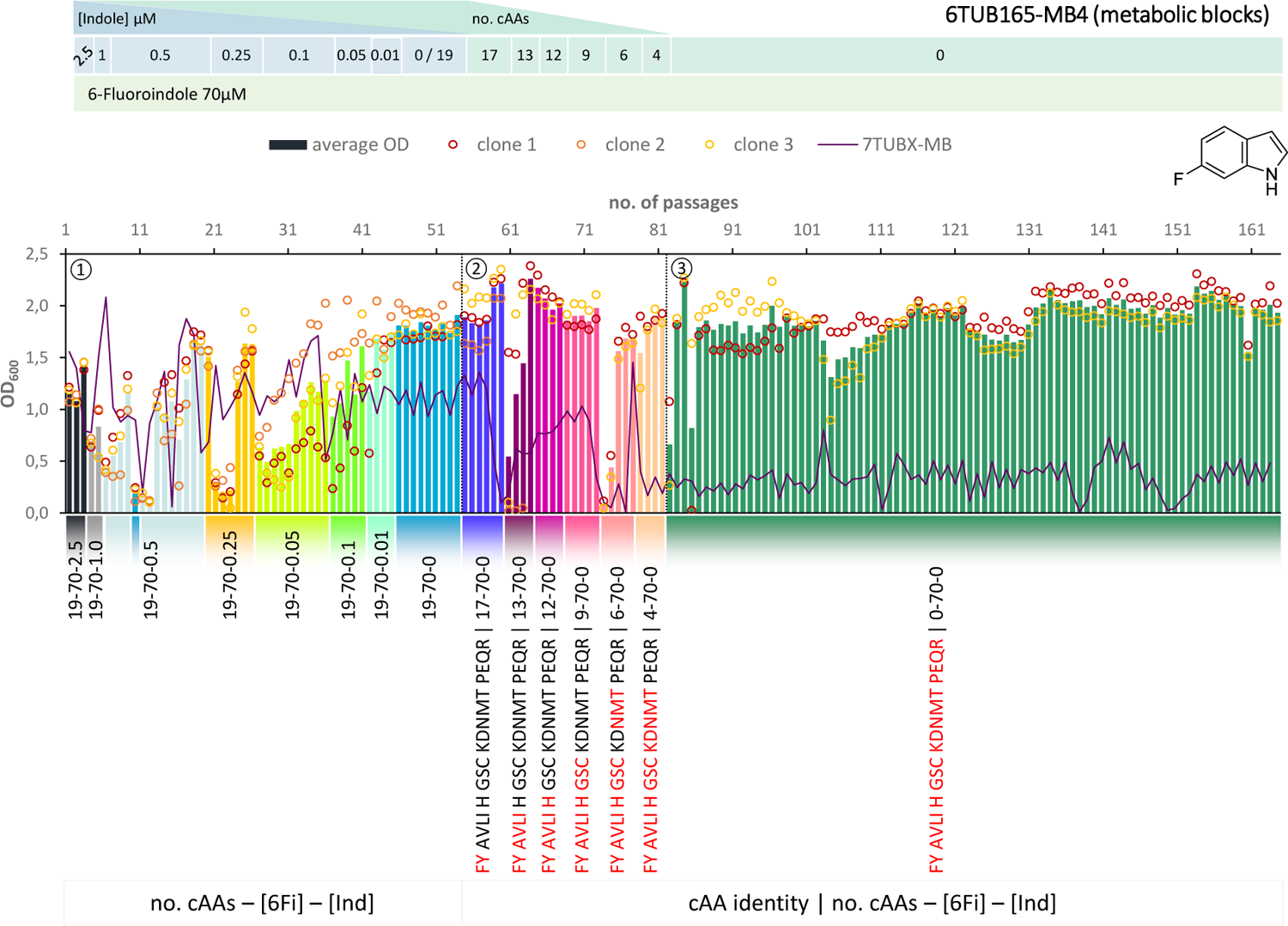
Cultivation scheme of the 6TUB165-MB4 lineage. Optical density (OD600) is plotted against the number of passages (reinoculation steps); average OD600 is shown as bars and individual clones are shown as circles. The corresponding 7TUBX-MB counterpart is depicted as purple line. ALE phases ①, ②, ③ and medium composition are illustrated; in the MB approach the amino acid composition is shown where red letters representing removed cAAs.

**Figure 6.**
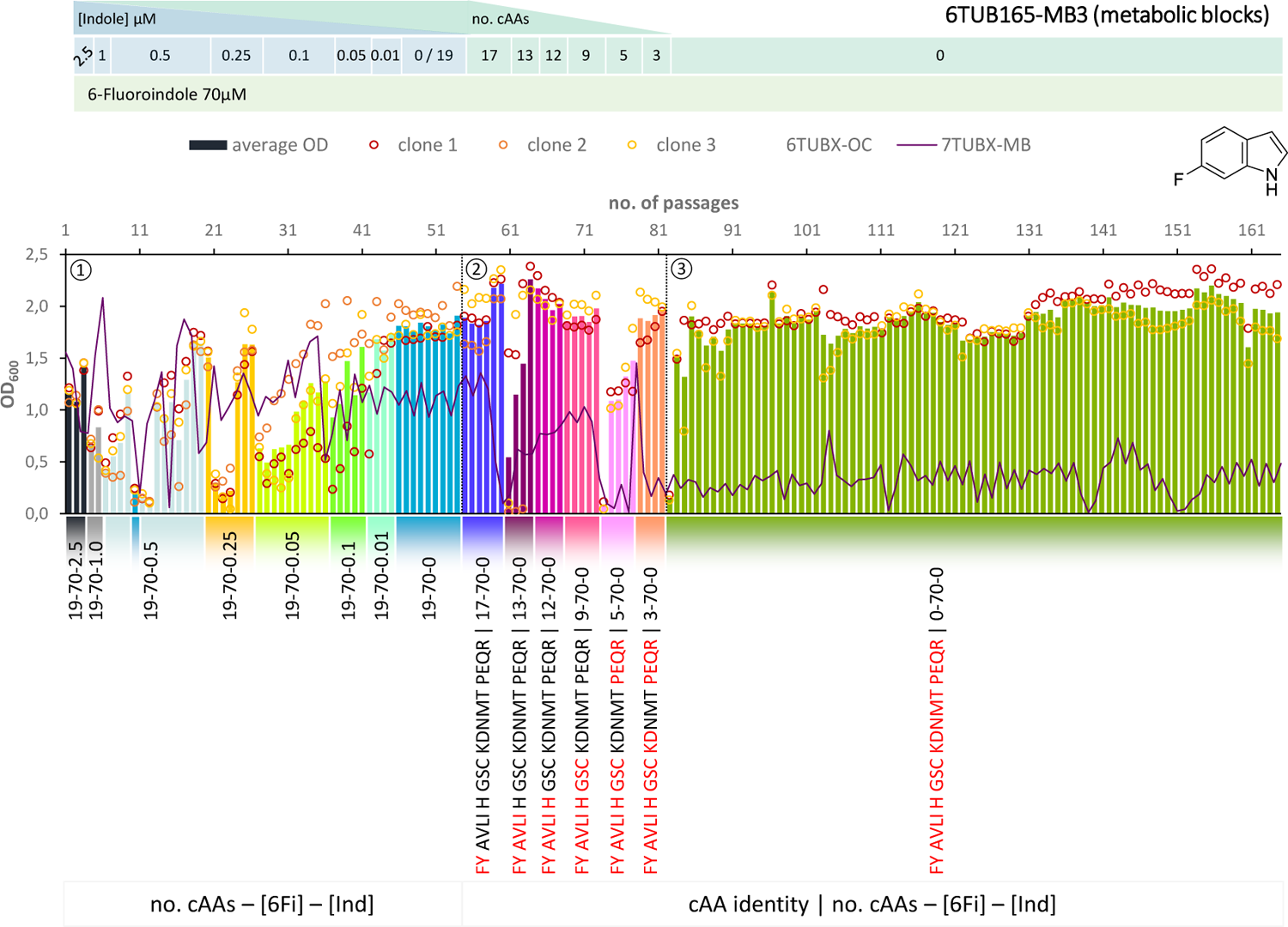
Cultivation scheme of the 6TUB165-MB3 lineage. Optical density (OD600) is plotted against the number of passages (reinoculation steps); average OD600 is shown as bars and individual clones are shown as circles. The corresponding 7TUBX-MB counterpart is depicted as purple line. ALE phases ①, ②, ③ and medium composition are illustrated; in the MB approach the amino acid composition is shown where red letters representing removed cAAs.

### Subpopulation screening

When screening for indole and fluoroindole compatibility, it was found that clones of both 6TUBX-MB lineages grew better in the presence of 6Fi than indole. Indeed, some clones were found to reject indole entirely when grown on solid medium (Figure 7A). This behavior was not observed in clones of the 6TUBX-OC lineage; thus, its cultivation was terminated after 128 passages (in contrast to the 165 passages of the 6TUBX-MB lineages).

**Figure 7.**
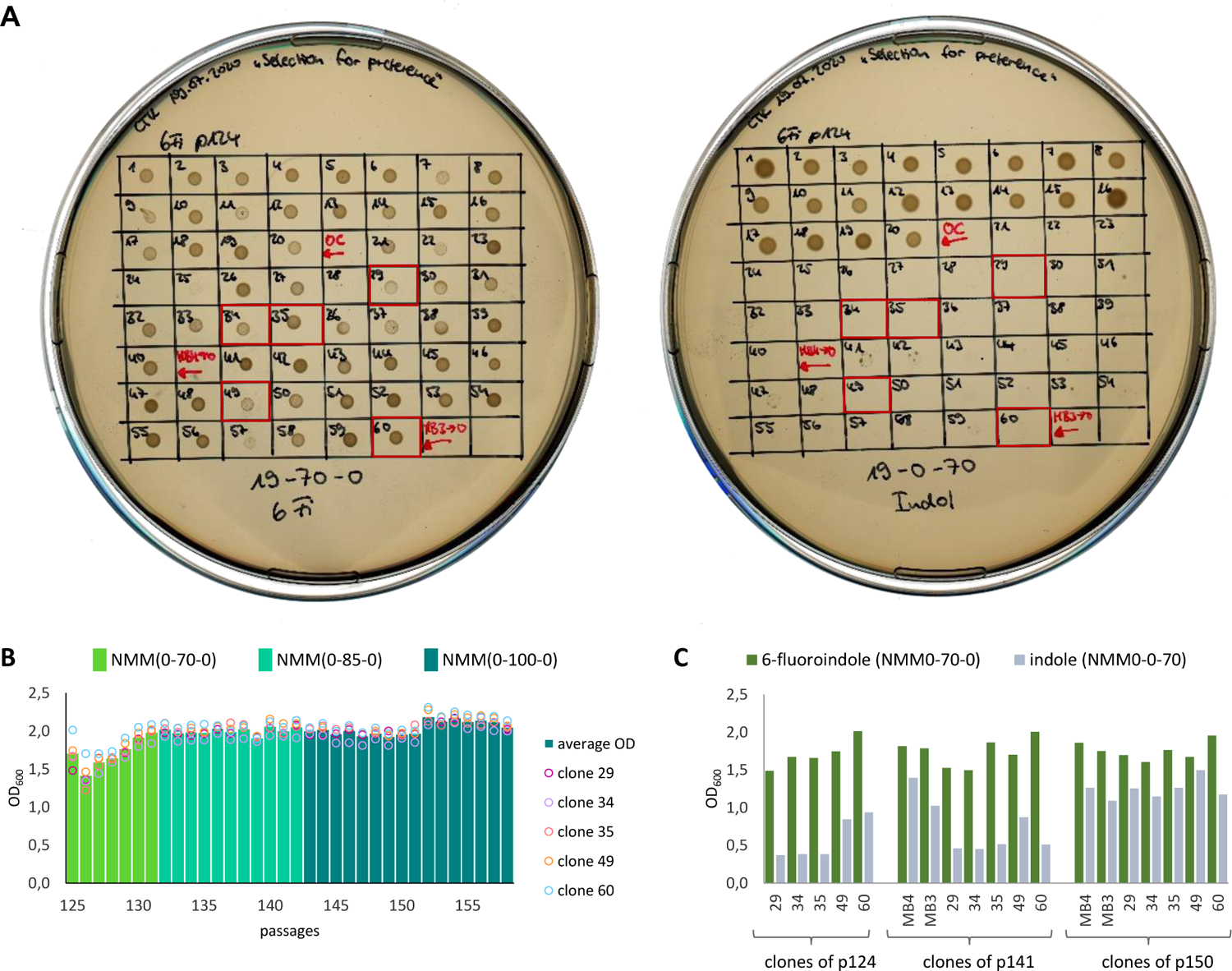
Subpopulation screening for a 6-fluoroindole preferring phenotype. A) Screening for 6Fi preferring clones by plate growth selection. Twenty individual clones of 6TUB124-OC, 6TUB124-MB4 and 6TUB124-MB3 were diluted and spotted on NMM19-70-0 (6Fi) and NMM19-0-70 (Ind) agar plates. The desired phenotype grew on the 6Fi supplemented plate but not on the indole-enriched plate. Promising clones 29, 34, 35 (from 6TUB124-MB4) and 49, 60 (from 6TUB124-M3) are outlined in red and are labeled with the suffix “SP”. B) Of the 60 clones tested, five promising clones were selected and subjected to an additional adaptation experiment in which the selective pressure (6Fi concentration) was gradually increased from 70 µM to 85 µM up to 100 µM. C) Screening of the subset of promising clones (29, 34, 35, 49, 60) and parent clones of 6TUBX-MB4 and 6TUBX-MB3 in liquid culture supplemented with either 6Fi or Ind (NMM0-70-0 or NMM0-0-70). Different time points of sub-ALE were chosen with passage 124 (70 µM 6Fi), passage 141 (85 µM 6Fi) and passage 150 (100 µM 6Fi). However, the parent lineages (MB4, MB3) were kept constantly on 70 µM 6Fi, therefore allowing to distinguish the effects of an increased 6Fi concentration from starting point (70 µM) on the progress.

The ultimate goal of our evolution experiments was to create organisms that are intransigently adapted to fluorine, i.e., to evolve cells that are not merely facultative F-Trp users. To this end, we selected five promising clones (29, 34, 35, 49, and 60) from the 6TUBX-MB strains and continued cultivation under enhanced selection pressure, by increasing the 6Fi concentration from 70 µM to 85 µM to 100 µM (Figure 7B). The resulting lineages were designated with the suffix “SP”, referring to subpopulation. Unfortunately, the previously indicated selectivity for 6Fi did not further increase even after 34 passages under these conditions. We tested these clones as well as the parent lineages (6TUBX-MB4 and 6TUBX-MB3) at different time points in liquid culture supplemented with either 6Fi or Ind (Figure 7C). When cultured in liquid medium, a remarkable difference is also observed in the optical densities achieved either with 6Fi or Ind, with a clear preference of the 6Fi-favoring phenotype. While this tendency is maintained in cells of passage 141, it is lost with passage 150. Most probably the increased selection pressure worked well up to 85 µM, but further increase to 100 µM did not produce the desired result of an unconditionally 6Fi-dependent phenotype. Further isolation and subpopulation screening did not yield any new clones, suggesting that this phenotype has been lost. Nevertheless, in-depth analyses (genetic, transcriptomic, metabolomic level) may shed light on possible mechanisms that caused the initial trend.

### Adaptive laboratory evolution experimental setup for 7-fluorindole (7Fi)

In contrast to 6Fi, tolerance towards 7Fi appears to be significantly more challenging as indicated by the low population sizes especially during the late adaptation process (Figures 8 and **9**). Already in phase 1, a high clonal variation in the acceptance to the growth conditions is observed, however, in sum similar high OD_600_ values were sustained as in the same phase of the 6Fi adaptation experiments, indicated by 6TUBX-MB reflection (green line) in Figures 8 and **9**. In phase 2 cell density decreased continuously in both lineages (OC and MB). We hypothesize that this resulted from the low-grade toxicity of 7Fi that became increasingly difficult for cells to compensate while undergoing cAA starvation. In case of the MB lineage, we alternatively tried the removal of different amino acid as well as performed replay experiments[46] with the same setup starting from passages 72 when growth was still acceptable to investigate if the sudden drop in the cell density is reproducible. However, all such attempts ended with drastic reductions in cell population.

**Figure 8.**
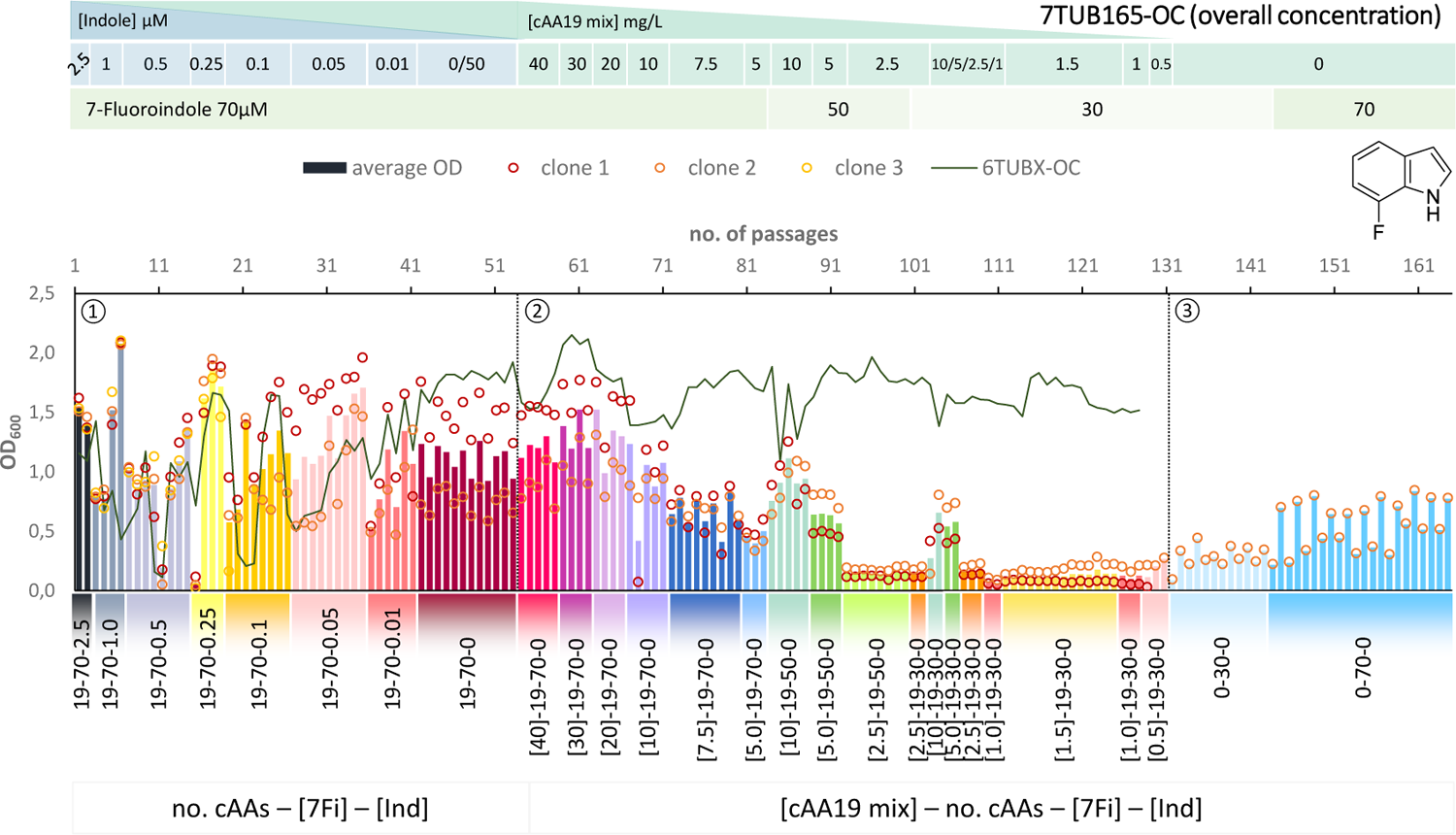
Cultivation scheme of the 7TUB165-OC lineage. Optical density (OD600) is plotted against the number of passages (reinoculation steps); average OD600 is shown as bars and individual clones are shown as circles. The corresponding 6TUBX-OC counterpart is depicted as green line. ALE phases ①, ②, ③ and medium composition are illustrated; in the OC approach the concentration (mg/L) of the cAA19 mix is indicated in squared brackets [cAA conc.]-a-b-c. Note the relaxed growth conditions by means of increased amino acid supply and reduced 7Fi concentration, depicted in the legends.

**Figure 9.**
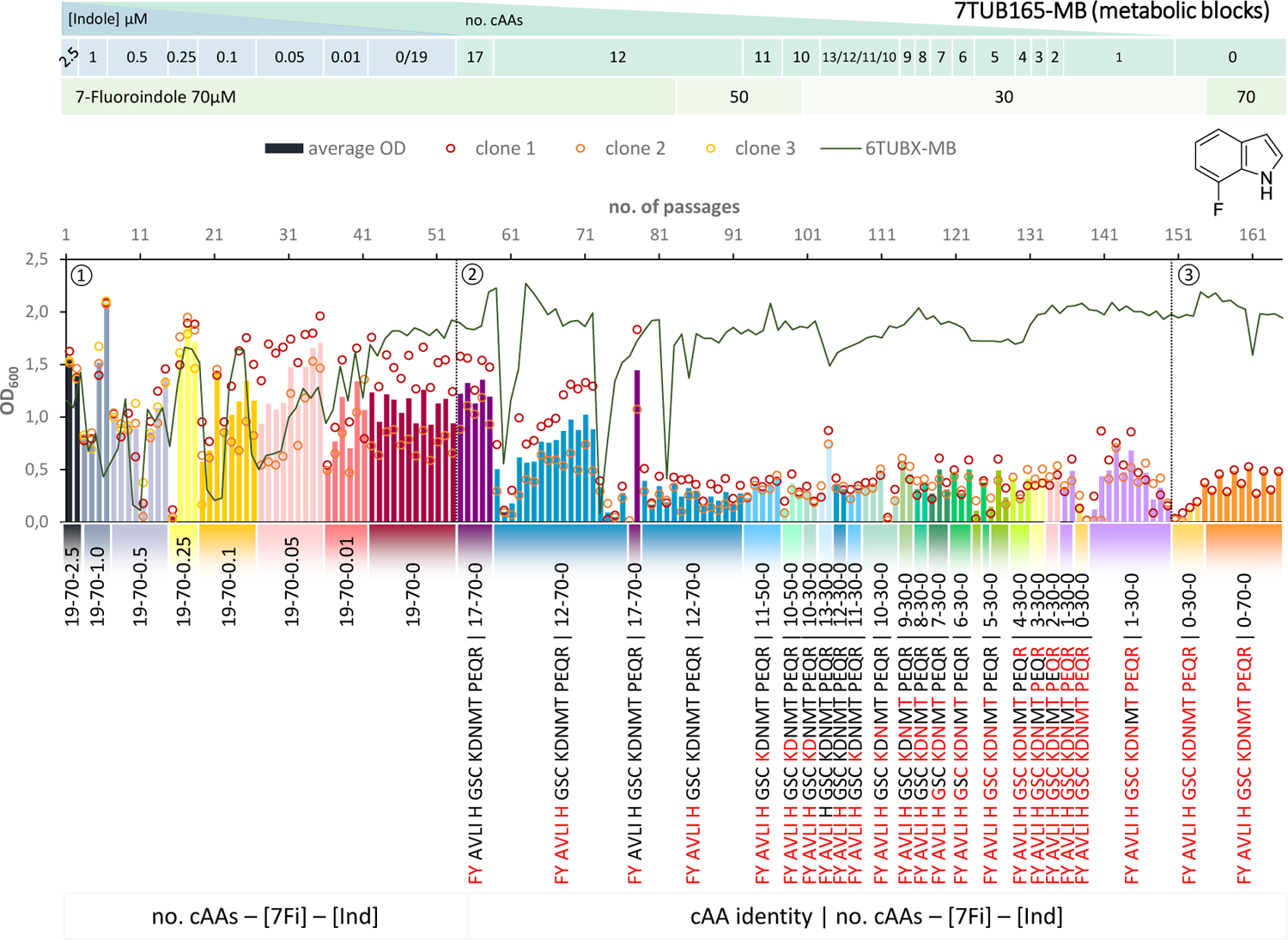
Cultivation scheme of the 7TUB165-MB lineage. Optical density (OD600) is plotted against the number of passages (reinoculation steps); average OD600 is shown as bars and individual clones are shown as circles. The corresponding 6TUBX-MB counterpart (mean of MB4 and MB3) is depicted as green line. ALE phases ①, ②, ③ and medium composition are illustrated; in the MB approach the amino acid composition is shown where red letters representing removed cAAs. Note the relaxed growth conditions by means of increased amino acid supply and reduced 7Fi concentration, depicted in the legends.

In order to rescue the cells, we reduced the concentration of 7Fi while temporarily increasing the cAA supply both starting with passage 84 (displayed in the legends of Figure 8 and Figure 9). Finally, the cells slowly recovered upon the relaxed selection pressure and the concentration of 7Fi could again raised to 70 µM. Still, this process demonstrated that cAAs had to be removed extremely conservatively (compared to 6Fi): in smaller steps in the OC approach and not at all as metabolic blocks in the MB approach, but rather removed on an individual basis. Even then not all clones survived the treatment, and the population sizes are considerably lower compared to the 6Fi adaptations. Taken together, this underlines the great challenge for *E. coli* to accept and adapt to 7Fi. Based on the described hurdles, a noteworthy difference between the OC and the MB approach is difficult to be determined, however, from an experimental point of view the OC approach was easier to implement.

### Growth characteristics of evolved strains

Growth characteristics serve as an excellent indicator for the success of adaptation and enable assessment of the phenotype and bacterial fitness of the adapted cells. To this end, we conducted extensive growth experiments of all strains in different media that reflect the conditions along the adaptation trajectory (**Figure 10B**, upper panel).

**Figure 10.**
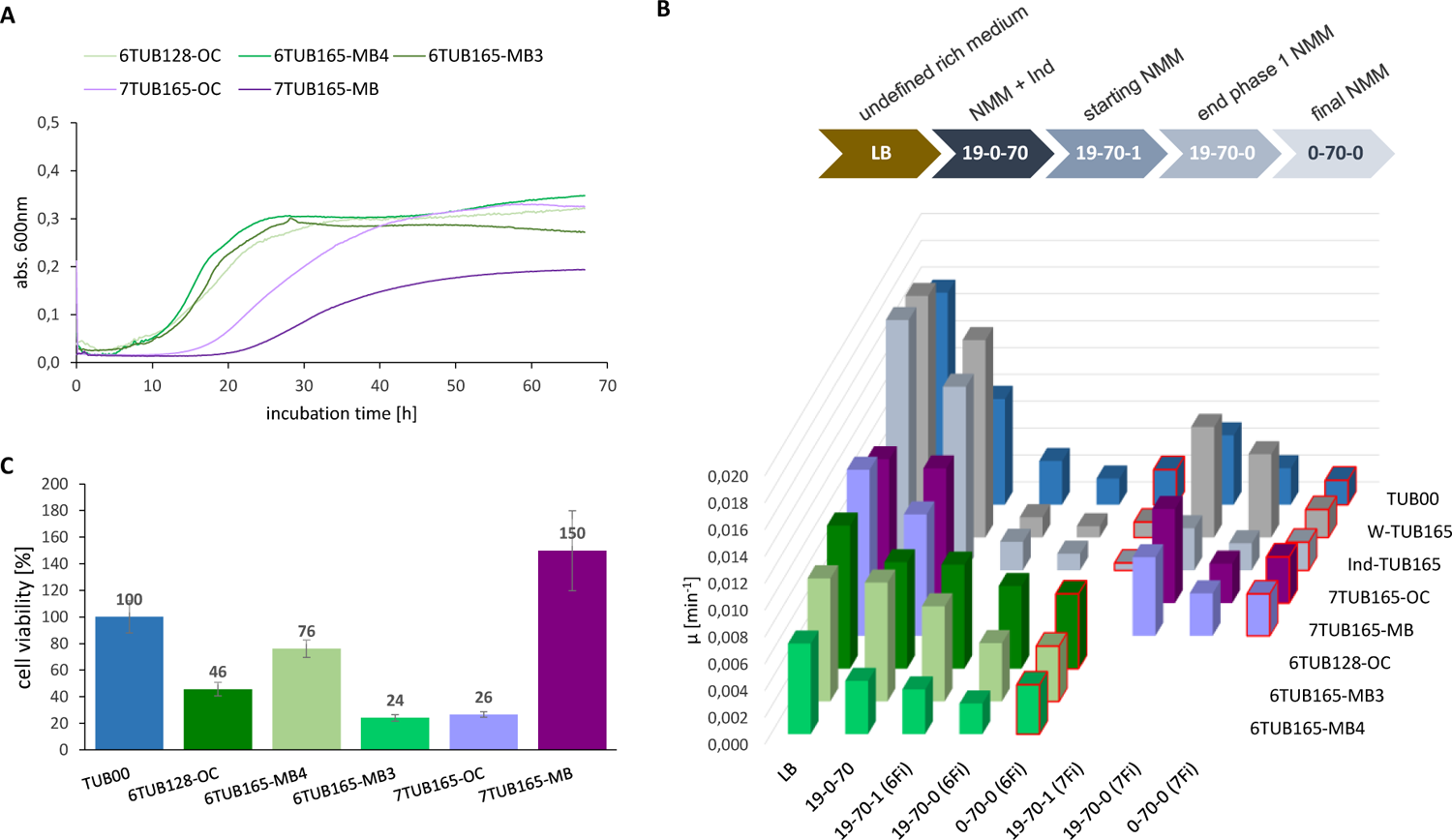
Growth characteristics of adapted strains (6TUB128-OC, 6TUB165-MB4, 6TUB165-MB3, 7TUB165-OC, 7TUB165-MB and W-TUB165, Ind-TUB165) in comparison to the ancestral TUB00. A) Entire growth curves of final isolates of 6Fi adapted (greenish) and 7Fi adapted (purple) strains measured by growth in their final medium composition NMM0-70-0 (70 µM 6Fi or 7Fi, respectively). B) The specific growth rate (µ) rate was calculated from the slope of exponential phase using the program GrowthRates;[47] values obtained in final medium composition are red framed and characteristics of the different growth media are depicted above. C) Cell viability was determined using CCK-8 assay (Cell Counting Kit-8); values of F-adapted strains were normalized to TUB00 (set 100 %). All measurements were performed at microscale level in triplicates using a plate reader.

Overall, the impression about the adaptation success gained from the cultivation schemes (**Figures 4, 5, 6, 8** and **9**) can also be seen in the growth curves (**Figure 10A**). The growth performance of 6Fi adapted cells is significantly better compared to the 7Fi adapted cells. The final isolates of the 6Fi adapted cells exhibit a very similar growth behavior in their final growth medium (NMM0-70-0), although 6TUB165-MB4 appears to be the best adapted lineage. In contrast, the growth behavior of cells adapted to 7Fi is clearly different. These cells exhibit a significant increased lag time and decreased growth rate, which is more pronounced in the 7TUB165-MB strain that also shows a lower maximum OD_600_ value.

Furthermore, **Figure 10B** provides an overview of the growth of all investigated strains along the adaptation course given by calculation of the specific growth rate[47] (for comprehensive view see detailed growth curves in Figures S10A-H). Generally, the growth correlates with the concentration of provided nutrients. Interestingly, the ancestral TUB00 tolerates 7Fi better than 6Fi, although based on the cultivation schemes (Figure 8, Figure 9) we have considered 7Fi to be more toxic. It is likely that 7Fi is metabolically more inert (less similar to indole), thus it affects the cellular metabolism less than 6Fi (which is more similar to indole and therefore confounds intracellular metabolism well). The growth performance of the positive controls in NMM19-0-70 approaches that in LB (see curves in Figures S10B-C), which is comprehensible because these strains were adapted to grow on minimal medium. On the other hand, the fluorinated indoles remain toxic or at least inhibiting to the positive controls.

Among the 6Fi adapted cells, the OC lineage exhibit the highest specific growth rate in all tested media. However, as also seen in the MB3 lineage, there is no further growth improvement observable after the transition from NMM19-70-0 to NMM0-70-0. Remarkably, the MB4 lineage displays such enhanced growth behavior. Although its growth rates are lower compared to both other lineages, in 6Fi supplemented NMM, the highest growth rate is observed in the final medium NMM0-70-0, which is only surpassed by growth in the ancestral medium. This finding is even more evident in the entire growth curves (see Figures S10D-F). This observation strongly indicates for approaching a fluoroindole preferring phenotype, which is also supported by the subpopulation screening results.

Both 7Fi-adapted strains show again a decreased growth rate along with reducing nutrient supply (cAAs, Ind). Indole supplemented media enables enhanced growth in absence of 7Fi (NMM19-0-70 > NMM19-70-1), which is a clear signal for an incomplete adaptation, as these cells still strongly prefer indole over 7Fi. However, the growth performance in 7Fi supplemented medium with and without cAAs addition (NMM19-70-0, NMM0-70-0) is equal, suggesting that removal of the canonical amino acids is well accepted, which seems surprising considering the challenging process that was required to adapt to these conditions.

Additionally, we completed the picture of bacterial fitness by determination of the cell viability using a CCK-8 assay (**Figure 10C**), which measures the bio-reduction of tetrazolium salts (WST-8).[48] The viability of all 6Fi adapted lineages correlates very well with the growth parameters, whereas among the 7Fi adapted strains there is a discrepancy found for 7TUB165-MB. This strain shows a cell viability of 150 % compared to TUB00, although the growth curves and parameters would suggest a considerably lower value.

### Susceptibility and tolerance of evolved strains to vancomycin

Beside obvious characteristics of growth behavior and viability we also investigated the membrane integrity of the adapted cells (Figure 11, Figure S11). In the course of our previous ALE study, the membranes of 4Fi and 5Fi adapted strains (4TUB93, 5TUB83) rearranged by an increased content of lipids containing unsaturated fatty acids, which at the same time increased their susceptibility towards antibiotics.[31] Therefore, we also tested the permeability of the cell membrane of TUB00 and the adapted strains by determination of the minimal inhibitory concentration (MIC) of vancomycin, an antibiotic inactive against gram-negative *E. coli* because the intrusion through the (intact) outer membrane is blocked.

**Figure 11.**
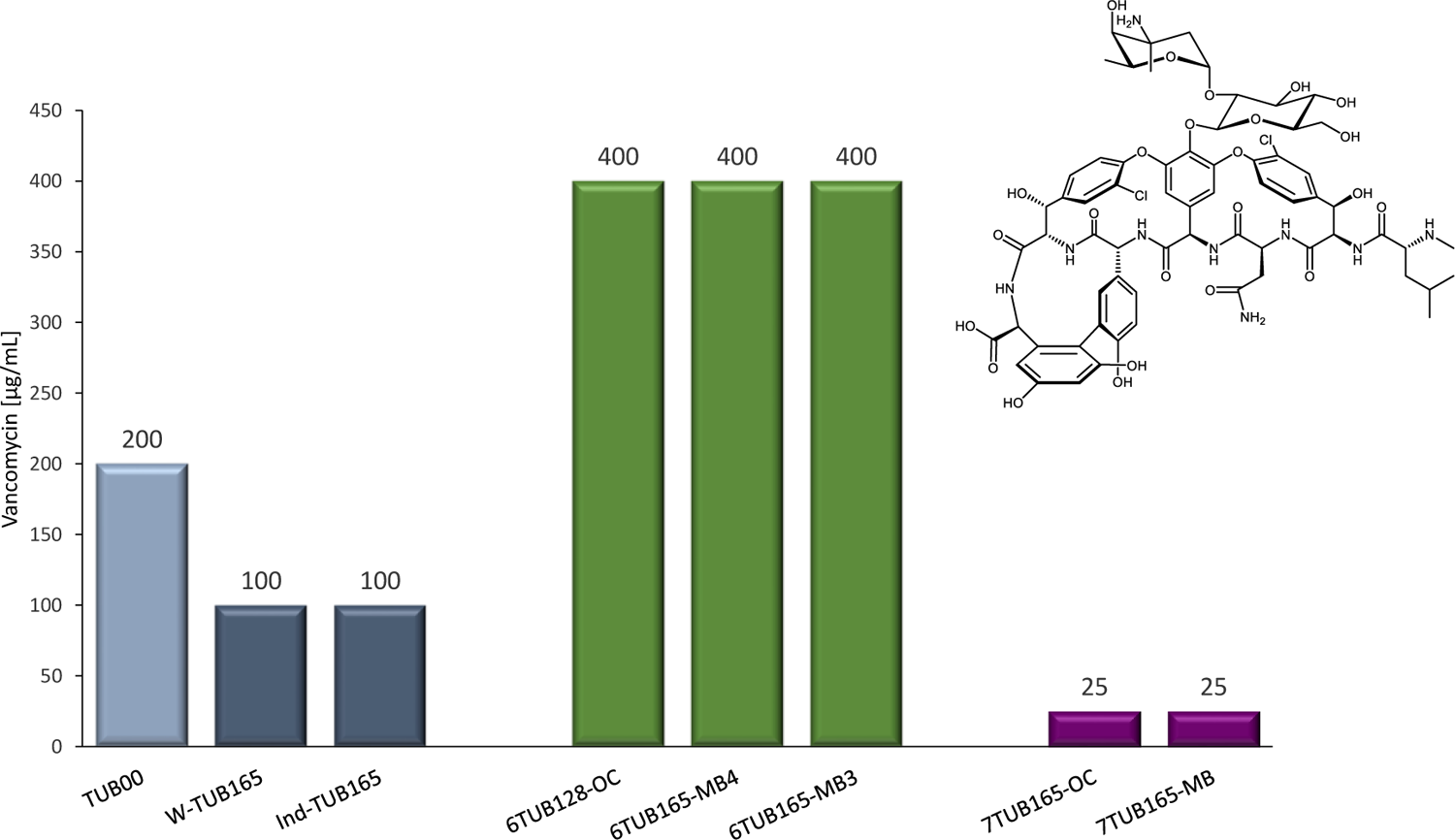
Investigation of cell membrane integrity by antibiotic susceptibility testing. The minimal inhibitory concentrations (MIC) of vancomycin are shown for the ancestral TUB00 (light grey), two positive controls W-TUB165 and Ind-TUB165 (grey), 6Fi adapted strains 6TUB128-OC, 6TUB165-MB4 and 6TUB165-MB3 (green) as well as 7Fi adapted strains 7TUB165-OC and 7TUB165-MB (purple). Cells were treated with a dilution series of vancomycin (0 - 400 µg/mL) and growth after 24 h incubation at 30 °C, 200 rpm is shown.

Compared to TUB00 (MIC 200 µg/mL) the positive controls show an increased vancomycin sensitivity (MIC 100 µg/mL). This might be a by temperature caused effect for one reason because membrane fluidity in general is influenced by temperature and for another because cold stress increases *E. coli*’s susceptibility to vancomycin.[49] As well as strains 4TUB93 and 5TUB83,[31] the 7Fi adapted cells also show a strongly increased vancomycin sensitivity (MIC 25 µg/mL), which can likely be attributed to a membrane rearrangement. Interestingly, the strains adapted to 6Fi exhibit the opposite effect being highly resistant to vancomycin treatment (MIC 400 µg/mL). One possible explanation is the accumulation of phosphatidic acid which is also a normal component of the membrane. But its increase is under suspicion to impede the entry of vancomycin and to confer resistance.[50] Of course, this theory has to be studied further for the adapted strains.

### Analytical evidence for proteome-wide replacement of Trp by 6FTrp or 7FTrp

Global substitution of Trp for FTrp was detected by protein nano-electrospray-ionization-mass-spectrometry coupled with liquid chromatography (nano-LC-ESI/MS). Trp-containing peptides were exclusively detected in TUB00 and both positive controls, while FTrp-containing peptides were only present in the isolates of adapted strains (see Figure S8).

Furthermore, the 6Fi and 7Fi adapted strains were tested for their robustness in driving recombinant protein expression.[51] Two different variants of the green fluorescent protein (GFP) model protein were used for heterologous expression; 1) the enhanced green fluorescent protein EGFP, which harbors one single Trp residue at position 57 in its primary sequence as well as 2) the enhanced cyan fluorescent protein ECFP, which contains also a Trp in the chromophore at position 66 (besides position 57) were used (cartoons in Figure 12).

**Figure 12.**
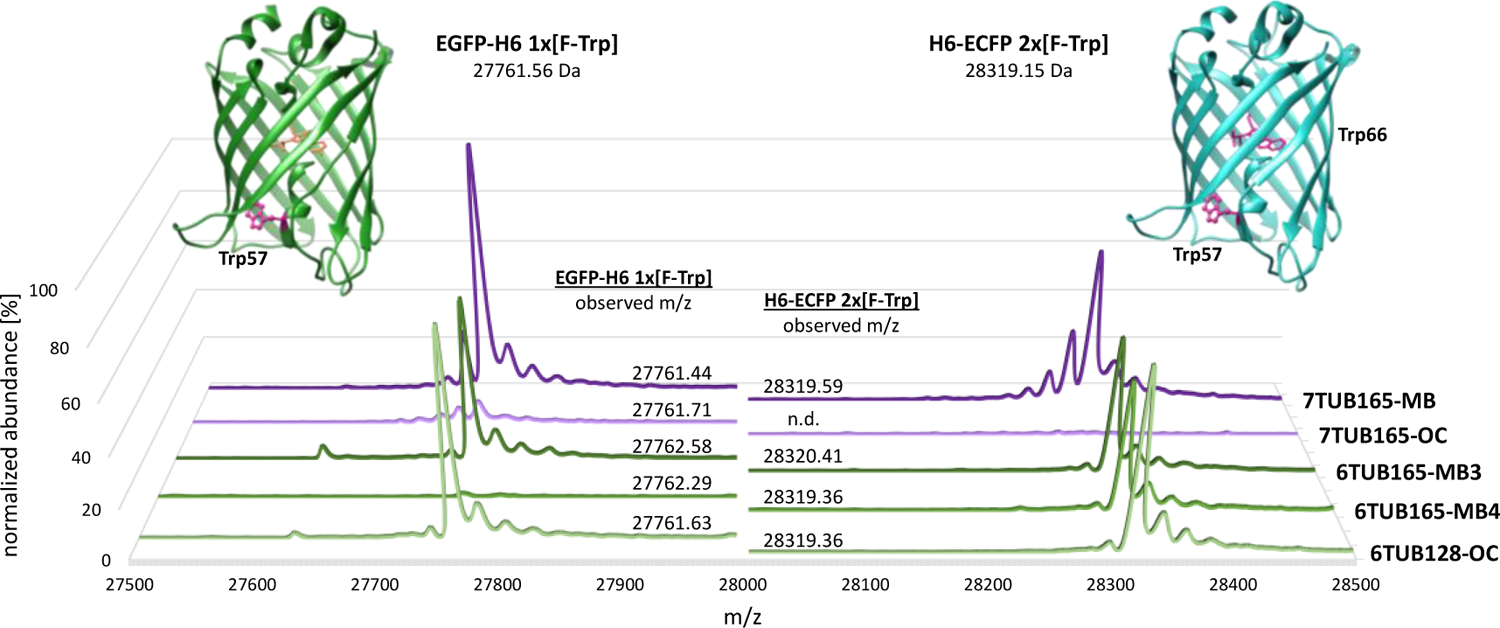
Deconvoluted mass spectra of C-terminal His-tagged enhanced green fluorescent protein (EGFP-H6, left) and N-terminal His-tagged enhanced cyan fluorescent protein (H6-ECFP, right). MS spectra of expression products of 6Fi adapted strains (6TUB128-OC, 6TUB165-MB4 and 6TUB165-MB3) are colored in green and respective spectra of proteins expressed with 7Fi adapted strains (7TUB165-OC and 7TUB165-MB) are colored in purple. The structures of EGFP (PDB ID 2Y0G)[52] and ECFP (PDB ID 2WSN)[53] from *Aequorea victoria* were drawn using UCSF Chimera;[54] Trp-positions are highlighted.

Cells of the final isolates of all adapted strains (6TUB128-OC, 6TUB165-MB4, 6TUB165-MB3 and 7TUB165-OC, 7TUB165-MB) were transformed with a plasmid bearing the sequence of either His-tagged EGFP (C-terminal EGFP-H6) or ECFP (N-terminal H6-ECFP). The recombinant protein expression was performed under the same conditions as used for adaptation, meaning cultivation in NMM0-70-0 containing either 6- or 7-fluoroindole at 30 °C; followed by isolation and purification using Ni-NTA affinity chromatography and analysis via LC-ESI-MS (see Figure S12 and Table S6).

Almost all adapted strains support recombinant expression of GFP variants in which all Trp residues are quantitatively replaced by fluorinated counterparts. Mass spectrometric analyses of 6FTrp- and 7FTrp-substituted EGFP and ECFP show homogeneous mass profiles associated with an expected mass shift of 18 Da (which correlates with a H-to-F exchange on the aromatic ring of the indole). The expected theoretical values for translated EGFP and ECFP after chromophore maturation[55] are consistent with the observed masses (Figure 12). However, the amount of 6FTrp-tagged EGFP expressed with 6TUB165-MB4 was very low; and the 7TUB165-OC strain was either able to produce only small amounts of 7FTrp-EGFP or was even unable to form 7FTrp-ECFP at all. Moreover, in addition to the dominant mass of the full-length protein, the spectra of the substituted EGFPs show a second signal assigned to a truncated product after processing of the initial methionine (mass: 27630 Da, Δm/z −131 Da). Due to the substitution of Trp66 in the chromophore triad of ECFP, a slight change in fluorescence profiles upon fluorination should be detectable.[56] The successful expression confirmed that in the adapted strains the protein synthesis was reprogrammed as Trp was completely replaced by 6- and 7-fluorotryptophan, respectively. We hypothesize that these findings can be extrapolated to the whole proteome and all Trp residues in the proteome of the adapted strains are completely replaced by 6- or 7-fluorotryptophan. In this way, the biological generation of fluorine-containing biomass becomes a reality.

## Discussion

The present adaptation experiments towards 6Fi and 7Fi convincingly demonstrate the reproducibility of our previous study on the adaptation towards 4Fi and 5Fi[31] as well as of the ALE process itself. Hence, we have established a reliable platform and methodologies for the experimental evolution of novel organisms with fluorine as a bioelement, created by mimicking natural selection in the laboratory.

In the course of the described adaptation process, the cells relinquished their dependence on canonical Trp and instead acquired the ability to use the fluorinated counterparts for all cellular processes e.g., consequent incorporation F-Trp into the proteome. This once again demonstrates the feasibility of assimilation of fluoroindoles and trophic reassignment of the UGG codon by fluorinated tryptophan. Based on the adaptation process itself and subsequent characterization experiments we observed considerable differences on the acceptance for 6Fi and 7Fi, where 6Fi is unambiguously better tolerated by *Escherichia coli* (Figure S6 and S10, Table 5). Overall, the 6Fi adapted lineages reach higher optical densities of about OD_600_ = 2 and higher specific growth rates, whereas the 7Fi adapted strains grew in the same incubation time to merely populations sizes of about OD_600_ = 0.5 – 0.8 and lower growth rates.

We expanded the efforts of the adaptation process by adding, for the first time, two positive controls long-term cultivated on either Trp or indole, as well as a new approach for amino acid removal. The positive controls will allow us to distinguish between general consequences of long-term propagation in minimal medium and those that are causative fluorine derived. They can also be consulted for our other Trp-based ALEs[30,31] since we conserved the same adaptation setup. Our novel OC approach for amino acid removal provides an excellent alternative for adaptation of challenging substrates and might be more suitable to apply for already stressed cells, as shown in case of 7-fluoroindole. In our opinion, this straightforward technique is more comfortable in laboratory handling but in addition we believe that the metabolic burden to the cells of biosynthesizing missing amino acids is reduced upon this method (compared to the MB approach). Nevertheless, both approaches led to the desired adaptation, notwithstanding the different cellular outcomes for instance that lineage 7TUB165-OC clearly exhibits better growth performance than 7TUB165-MB, but that in terms of recombinant protein expression capability an opposite tendency is observed. Therefore, so we do not want to single out one approach over the other.

Contrary to 4-, 5- and 6-fluoroindole/Trp, the Ind/Trp isomer fluorinated at position 7 is so far rather underrepresented in studies on biosynthetically incorporation and only a few have investigated the differences of all four isomers in specific proteins in direct comparison.[56–59] Why these isomers have not made much of an appearance remains speculative, however, the chemical synthesis and use of both 7Fi[60] and 7FTrp[61,62] was already described many decades ago.

However, the proteome-wide insertion of 7-fluorotryptophan (by cellular conversion of 7-fluoroindole) described here represents the first study of its kind involving this non-canonical amino acid. Our findings indicate that adaptation to 7Fi is highly challenging for *E. coli*, evident through the intricate nature of the adaptation process and the compromised growth performance of the cells. Exciting opportunities lie ahead in future studies to unravel the molecular factors responsible for these difficulties, which likely stem from the chemistry and structure of this analogue and its impact on cellular metabolism. Potential areas of investigation include its effects on metabolic pathways and the influence on protein folding quality for proteins containing 7-fluorotryptophan.

The adaptation experiments towards usage of 6Fi, on the other hand, showed an astonishing readiness of the cells for this isomer. Here, the most notable outcome was the emergence of a subpopulation within the MB lineages that had started to become nearly independent of indole and even prefer 6Fi. Equally astonishing was the finding that all 6Fi adapted lineages showed an increased resistance to vancomycin, which is unique among all other 4-, 5-,[31] and 7Fi adapted strains.

Proteome-wide substitution of Trp by FTrp was demonstrated for all fluoroindole-based ALEs. However, although *E. coli* had acquired the competence to use the fluorinated substrates as integrated metabolic intermediates, the adapted strains remained (still) facultative fluorotryptophan/tryptophan users. In addition, they appear to be exclusively specified to their adapted indole analogue since no growth advantages of the adapted cells over their other mono-fluorinated analogues could be observed, that is 6TUBX strains on 4-, 5- or 7Fi or else 7TUBX on 4-, 5- or 6Fi.[27,29]

Furthermore, our fluorine adapted strains are appropriate expression hosts to readily produce FTrp substituted proteins and peptides; thus, opens up opportunity to go beyond and overcome limitations of automated solid-phase peptide synthesis (SSPS) by harnessing the ribosomal synthesis of adapted cells.[63] However, the application of classical molecular biology methods such as amber stop codon suppression (SCS) and selective pressure incorporation (SPI)[64] also benefit from the abolished dependence on canonical amino acids of the adapted strains, allowing improved and effective synthesis of fully fluorine-labeled proteins.

Although largely ignored by nature, a life based on fluorine is both an interesting concept and an absolutely conceivable scenario.[65] Fluorine-containing building blocks have been extensively used to investigate and modify proteins and their interactions.[66–68] However, the adaptation of living organisms represents a step forward in exploring and understanding the consequences of global fluorination. At this point, we would like to highlight the outstanding achievements of a cell to cope with the intricate and myriad consequences associated with the exchange of a hydrogen against a fluorine atom in an essential building block (after all more than 20000 Trp positions only in the proteome).[44] Ultimately, this affects the complex interplay of cellular processes that include not only protein interactions through incorporated F-Trp such as π-π stacking, hydrogen bonding, and cation-π interactions but also all metabolic processes to which Trp or downstream metabolites contribute to.

Our adaptation experiments using fluorinated Trp precursor will contribute to a better understanding of the mechanism required to enable life with this extraordinary element. With this study, we completed the set of *Escherichia coli* bacteria adapted to all mono-fluorinated indoles (4-, 5-, 6- and 7Fi) and proofed them to be suitable metabolic substrates for the assembly of fluorinated proteomes. Based on this achievement we can now investigate the influence of the position of the F-substituent on cellular adaptability correlating our results with the physicochemical properties of the constitutional isomers, such as polarity and lipophilicity.[31] Even a cursory comparison of the cultivation schemes of all four substrates shows parallels in the adaptation behavior of 4Fi and 6Fi, which are well tolerated, while 5Fi and 7Fi present adaptation challenges.

This study with its focus on improving protocols and procedures provides reliable empirical determination of parameters for any ALE experiment on microbial adaptation to non-canonical amino acids. We strongly believe that the here established procedures will become one of the main routes in the engineering of synthetic cells with a chemically modified make-up of life. In future direction, we will dive into the different biological information level (DNA, RNA, proteins and metabolites) by means of multi-OMIC experiments to gain an understanding of the processes underlying adaptation and to compare them with earlier studies in order to unravel a rational behind the adaptation towards xenobiotics and fluorine in particular. Enlightening the responsible mechanisms will push advancements in scientific fields such as synthetic biology and biotechnology, but also across the bord of environmental concerns such as bioremediation and biocontainment.[69]

## Authors contribution

C. T.-K. wrote the manuscript and performed experiments. C. T.-K., N. B. and B. K. conceived the project and designed the experimental setup, L. A. conducted the proteomic analyses and A. B. contributed to the writing of the manuscript at various stages.

## Data and Materials availability

Materials and methods, as well as additional data, are available in the separate SI file.

## References

[1] Harper, D. B. and O’Hagan, D. The fluorinated natural products. Nat. Prod. Rep. 11, 123–133 (1994).

[2] Carvalho, M. F. and Oliveira, R. S. Natural production of fluorinated compounds and biotechnological prospects of the fluorinase enzyme. Crit. Rev. Biotechnol. 37, 880–897 (2017).

[3] Sanada, M., Miyano, T. and Iwadare, S. Biosynthesis of fluorothreonine and fluoroacetic acid by the thienamycin producer, Streptomyces cattleya. J. Antibiot. (Tokyo). 39, 259–265 (1986).

[4] O’Hagan, D., Schaffrath, C., Cobb, S. L., Hamilton, J. T. G. and Murphy, C. D. Biosynthesis of an organofluorine molecule. Nature 416, 279–279 (2002).

[5] Deng, H., O’Hagan, D. and Schaffrath, C. Fluorometabolite biosynthesis and the fluorinase from Streptomyces cattleya. Nat. Prod. Rep. 21, 773–784 (2004).

[6] Markakis, K., Lowe, P. T., Davison-Gates, L., O’Hagan, D., Rosser, S. J. and Elfick, A. An Engineered E. coli Strain for Direct in Vivo Fluorination. ChemBioChem 21, 1856–1860 (2020).

[7] Eustáquio, A. S., O’Hagan, D. and Moore, B. S. Engineering fluorometabolite production: Fluorinase expression in salinispora tropica yields fluorosalinosporamide. J. Nat. Prod. 73, 378–382 (2010).

[8] Calero, P., Volke, D. C., Lowe, P. T., Gotfredsen, C. H., O’Hagan, D. and Nikel, P. I. A fluoride-responsive genetic circuit enables in vivo biofluorination in engineered Pseudomonas putida. Nat. Commun. 11, (2020).

[9] Dall’Angelo, S., Bandaranayaka, N., Windhorst, A. D., Vugts, D. J., van der Born, D., Onega, M., Schweiger, L. F., Zanda, M. and O’Hagan, D. Tumour imaging by Positron Emission Tomography using fluorinase generated 5-[18F]fluoro-5-deoxyribose as a novel tracer. Nucl. Med. Biol. 40, 464–470 (2013).

[10] Merkel, L. and Budisa, N. Organic fluorine as a polypeptide building element: in vivo expression of fluorinated peptides, proteins and proteomes. Org. Biomol. Chem. 10, 7241–7261 (2012).

[11] O’Hagan, D. and Deng, H. Enzymatic fluorination and biotechnological developments of the fluorinase. Chem. Rev. 115, 634–649 (2015).

[12] Gribble, G. W. The diversity of naturally produced organohalogens. Chemosphere 52, 289–297 (2003).

[13] Gribble, G. W. A recent survey of naturally occurring organohalogen compounds. Environ. Chem. 12, 396–405 (2015).

14. Neumann, C. S., Fujimori, D. G. and Walsh, C. T. Halogenation Strategies In Natural Product Biosynthesis. Chem. Biol. 15, 99–109 (2008).

[15] Gupta, G. N., Srivastava, S., Khare, S. K. and Prakash, V. Extremophiles: An Overview of Microorganism from Extreme Environment. Int. J. Agric. Enviroment Biotechnol. 7, 371–379 (2014).

[16] Merino, N., Aronson, H. S., Bojanova, D. P., Feyhl-Buska, J., Wong, M. L., Zhang, S. and Giovannelli, D. Living at the extremes: Extremophiles and the limits of life in a planetary context. Front. Microbiol. 10, 780 (2019).

[17] Yoshida, S., Hiraga, K., Takehana, T., Taniguchi, I., Yamaji, H., Maeda, Y., Toyohara, K., Miyamoto, K., Kimura, Y. and Oda, K. A bacterium that degrades and assimilates poly(ethylene terephthalate). Science (80-.). 353, 759 (2016).

[18] Sangwan, N., Verma, H., Kumar, R., Negi, V., Lax, S., Khurana, P., Khurana, J. P., Gilbert, J. A. and Lal, R. Reconstructing an ancestral genotype of two hexachlorocyclohexane-degrading Sphingobium species using metagenomic sequence data. ISME J. 8, 398–408 (2014).

[19] Wang, Z., Dewitt, J. C., Higgins, C. P. and Cousins, I. T. A Never-Ending Story of Per- and Polyfluoroalkyl Substances (PFASs)? Environ. Sci. Technol. 51, 2508–2518 (2017).

[20] Dragosits, M. and Mattanovich, D. Adaptive laboratory evolution - principles and applications in industrial biotechnology. Microb. Cell Fact. 12, 1–17 (2013).

[21] Sandberg, T. E., Salazar, M. J., Weng, L. L., Palsson, B. O. and Feist, A. M. The emergence of adaptive laboratory evolution as an efficient tool for biological discovery and industrial biotechnology. Metab. Eng. 56, 1–16 (2019).

[22] Connolly, J. P. R., Roe, A. J. and O’Boyle, N. Prokaryotic life finds a way: insights from evolutionary experimentation in bacteria. Crit. Rev. Microbiol. 47, 126–140 (2021).

[23] Wang, G., Li, Q., Zhang, Z., Yin, X., Wang, B. and Yan, X. Recent progress in adaptive laboratory evolution of industrial microorganisms. J. Ind. Microbiol. Biotechnol. kuac023, (2022).

[24] Mavrommati, M., Daskalaki, A., Papanikolaou, S. and Aggelis, G. Adaptive laboratory evolution principles and applications in industrial biotechnology. Biotechnol. Adv. 54, 107795 (2022).

[25] Barik, S. The uniqueness of tryptophan in biology: Properties, metabolism, interactions and localization in proteins. Int. J. Mol. Sci. 21, 8776 (2020).

[26] Tze Fei Wong, J. Membership mutation of the genetic code: Loss of fitness by tryptophan. Proc. Natl. Acad. Sci. U. S. A. 80, 6303–6306 (1983).

[27] Mat, W. K., Xue, H. and Wong, J. T. F. Genetic code mutations: The breaking of a three billion year invariance. PLoS One 5, 1–7 (2010).

[28] Yu, A. C. S., Yim, A. K. Y., Mat, W. K., Tong, A. H. Y., Lok, S., Xue, H., Tsui, S. K. W., Wong, J. T. F. and Chan, T. F. Mutations enabling displacement of tryptophan by 4-fluorotryptophan as a canonical amino acid of the genetic code. Genome Biol. Evol. 6, 629–641 (2014).

[29] Bacher, J. M. and Ellington, A. D. Selection and Characterization of Escherichia coli Variants Capable of Growth on an Otherwise Toxic Tryptophan Analogue. J. Bacteriol. 183, 5414–5425 (2001).

30. Hoesl, M. G., Oehm, S., Durkin, P., Darmon, E., Peil, L., Aerni, H. R., Rappsilber, J., Rinehart, J., Leach, D., Söll, D. and Budisa, N. Chemical Evolution of a Bacterial Proteome. Angew. Chemie - Int. Ed. 54, 10030– 10034 (2015).

[31] Agostini, F., Sinn, L., Petras, D., Schipp, C. J., Kubyshkin, V., Berger, A. A., Dorrestein, P. C., Rappsilber, J., Budisa, N. and Koksch, B. Multiomics Analysis Provides Insight into the Laboratory Evolution of Escherichia coli toward the Metabolic Usage of Fluorinated Indoles. ACS Cent. Sci. 7, 81–92 (2021).

32. Budisa, N. Prolegomena to future experimental efforts on genetic code engineering by expanding its amino acid repertoire. Angew. Chemie - Int. Ed. 43, 6426–6463 (2004).

[33] Budisa, N. and Pal, P. P. Designing novel spectral classes of proteins with a tryptophan-expanded genetic code. Biol. Chem. 385, 893–904 (2004).

[34] Tack, D. S., Ellefson, J. W., Thyer, R., Wang, B., Gollihar, J., Forster, M. T. and Ellington, A. D. Addicting diverse bacteria to a noncanonical amino acid. Nat. Chem. Biol. 12, 138–140 (2016).

[35] Watkins-Dulaney, E., Straathof, S. and Arnold, F. Tryptophan Synthase: Biocatalyst Extraordinaire. ChemBioChem 22, 5–16 (2021).

[36] Wilcox, M. The Enzymatic Synthesis of L-Tryptophan Analogues. Anal. Biochem. 440, 436–440 (1974).

[37] Goss, R. J. M. and Newill, P. L. A. A convenient enzymatic synthesis of L-halotryptophans. Chem. Commun. 4924–4925 (2006). doi:10.1039/b611929h

[38] Azim, M. K. and Budisa, N. Docking of tryptophan analogs to trytophanyl-tRNA synthetase: Implications for non-canonical amino acid incorporations. Biol. Chem. 389, 1173–1182 (2008).

[39] Blattner, F. R., Plunkett III, G., Bloch, C. A., Perna, N. T., Burland, V., Riley, M., Collado-Vides, J., Glasner, J. D., Rode, C. K., Mayhew, G. F., Gregor, J., Davis, N. W., Kirkpatrick, H. A., Goeden, M. A., Rose, D. J., Mau, B. and Shao, Y. The complete genome sequence of Escherichia coli K-12. Science (80-.). 277, 1453–1462 (1997).

[40] Vasi, F., Travisano, M. and Lenski, R. E. Long-term experimental evolution in Escherichia coli. II. Changes in life-history traits during adaptation to a seasonal environment. Am. Nat. 144, 432–456 (1994).

[41] Gresham, D. and Dunham, M. J. The enduring utility of continuous culturing in experimental evolution. Genomics 104, 399–405 (2014).

[42] LaCroix, R. A., Palsson, B. O. and Feist, A. M. A Model for Designing Adaptive Laboratory Evolution Experiments. Appl. Environ. Microbiol. 83, 1–14 (2017).

[43] Akashi, H. and Gojobori, T. Metabolic efficiency and amino acid composition in the proteomes of Escherichia coli and Bacillus subtilis. Proc. Natl. Acad. Sci. U. S. A. 99, 3695–3700 (2002).

[44] Tolle, I., Oehm, S., Hoesl, M. G., Treiber-Kleinke, C., Peil, L., Bozukova, M., Albers, S., Bukari, A.-R. A., Semmler, T., Rappsilber, J., Ignatova, Z., Gerstein, A. and Budisa, N. Evolving a mitigation of the stress response pathway to change the basic chemistry of life. *Front*. Synth. Biol. 1, 1248065 (2023).

[45] Wong, H. E., Huang, C.-J. and Zhang, Z. Amino Acid Misincorporation Propensities Revealed through Systematic Amino Acid Starvation. Biochemistry 57, 6767–6779 (2018).

[46] Blount, Z. D., Lenski, R. E. and Losos, J. B. Contingency and determinism in evolution: Replaying life’s tape. Science (80-.). 362, 655 (2018).

[47] Hall, B. G., Acar, H., Nandipati, A. and Barlow, M. Growth rates made easy. Mol. Biol. Evol. 31, 232–238 (2013).

[48] Yang, X., Zhong, Y., Wang, D. and Lu, Z. A simple colorimetric method for viable bacteria detection based on cell counting Kit-8. Anal. Methods 13, 5211–5215 (2021).

[49] Stokes, J. M., French, S., Ovchinnikova, O. G., Bouwman, C., Whitfield, C. and Brown, E. D. Cold Stress Makes Escherichia coli Susceptible to Glycopeptide Antibiotics by Altering Outer Membrane Integrity. Cell Chem. Biol. 23, 267–277 (2016).

[50] Sutterlin, H. A., Zhang, S. and Silhavya, T. J. Accumulation of phosphatidic acid increases vancomycin resistance in Escherichia coli. J. Bacteriol. 196, 3214–3220 (2014).

[51] Kuthning, A., Durkin, P., Oehm, S., Hoesl, M. G., Budisa, N. and Süssmuth, R. D. Towards Biocontained Cell Factories: An Evolutionarily Adapted Escherichia coli Strain Produces a New-to-nature Bioactive Lantibiotic Containing Thienopyrrole-Alanine. *Nat*. Sci. Rep. 33447 (2016). doi:10.1038/srep33447

[52] Royant, A. and Noirclerc-Savoye, M. Stabilizing role of glutamic acid 222 in the structure of Enhanced Green Fluorescent Protein. J. Struct. Biol. 174, 385–390 (2011).

[53] Lelimousin, M., Noirclerc-Savoye, M., Lazareno-Saez, C., Paetzold, B., Le Vot, S., Chazal, R., Macheboeuf, P., Field, M. J., Bourgeois, D. and Royant, A. Intrinsic dynamics in ECFP and cerulean control fluorescence quantum yield. Biochemistry 48, 10038–10046 (2009).

[54] Pettersen, E. F., Goddard, T. D., Huang, C. C., Couch, G. S., Greenblatt, D. M., Meng, E. C. and Ferrin, T. E. UCSF Chimera - A visualization system for exploratory research and analysis. J. Comput. Chem. 25, 1605–1612 (2004).

[55] Lundqvist, M., Thalén, N., Volk, A.-L., Hansen, H. G., von Otter, E., Nygren, P.-Å., Uhlen, M. and Rockberg, J. Chromophore pre-maturation for improved speed and sensitivity of split-GFP monitoring of protein secretion. Sci. Rep. 9, 310 (2019).

[56] Budisa, N., Pal, P. P., Alefelder, S., Birle, P., Krywcun, T., Rubini, M., Wenger, W., Bae, J. H. and Steiner, T. Probing the role of tryptophans in Aequorea victoria green fluorescent proteins with an expanded genetic code. Biol. Chem. 385, 191–202 (2004).

[57] Kenward, C., Shin, K. and Rainey, J. K. Mixed Fluorotryptophan Substitutions at the Same Residue Expand the Versatility of 19F Protein NMR Spectroscopy. Chem. Eur. J. 24, 3391–3396 (2018).

[58] Tobola, F., Lelimousin, M., Varrot, A., Gillon, E., Darnhofer, B., Blixt, O., Birner-Gruenberger, R., Imberty, A. and Wiltschi, B. Effect of Noncanonical Amino Acids on Protein-Carbohydrate Interactions: Structure, Dynamics, and Carbohydrate Affinity of a Lectin Engineered with Fluorinated Tryptophan Analogs. ACS Chem. Biol. 13, 2211–2219 (2018).

[59] Zlatopolskiy, B. D., Zischler, J., Schäfer, D., Urusova, E. A., Guliyev, M., Bannykh, O., Endepols, H. and Neumaier, B. Discovery of 7-[18F]Fluorotryptophan as a Novel Positron Emission Tomography (PET) Probe for the Visualization of Tryptophan Metabolism in Vivo. J. Med. Chem. 61, 189–206 (2018).

[60] Allen, F. L., Brunton, J. C. and Suschitzky, H. Heterocyclic Fluorine Compomds. Part II.* Bz-Monofluoroindoles. J. Chem. Soc. 1283–1286 (1955).

[61] Nevinsky, G. A., Favorova, O. O., Lavrik, O. I., Petrova, T. D., Kochkina, L. L. and Savchenko, T. I. Fluorinated tryptophans as substrates and inhibitors of the ATP-[32P]PPi exchange reaction catalyzed by tryptophanyl tRNA synthetase. FEBS Lett. 43, 135–138 (1974).

[62] Lee, M. and Phillips, R. S. Synthesis and resolution of 7-fluorotryptophans. *Bioorganic Med*. Chem. Lett. 1, 477–480 (1991).

[63] Völler, J. S., Dulic, M., Gerling-Driessen, U. I. M., Biava, H., Baumann, T., Budisa, N., Gruic-Sovulj, I. and Koksch, B. Discovery and Investigation of Natural Editing Function against Artificial Amino Acids in Protein Translation. ACS Cent. Sci. 3, 73–80 (2017).

[64] Hoesl, M. G. and Budisa, N. Recent advances in genetic code engineering in Escherichia coli. Curr. Opin. Biotechnol. 23, 751–757 (2012).

[65] Budisa, N., Kubyshkin, V. and Schulze-Makuch, D. Fluorine-Rich Planetary Environments as Possible Habitats for Life. Life 4, 374–385 (2014).

[66] Berger, A. A., Völler, J. S., Budisa, N. and Koksch, B. Deciphering the Fluorine Code - The Many Hats Fluorine Wears in a Protein Environment. Acc. Chem. Res. 50, 2093–2103 (2017).

[67] Monkovic, J. M., Gibson, H., Sun, J. W. and Montclare, J. K. Fluorinated Protein and Peptide Materials for Biomedical Applications. Pharmaceuticals 15, (2022).

[68] Marsh, E. N. G. and Suzuki, Y. Using 19F NMR to probe biological interactions of proteins and peptides. ACS Chem. Biol. 9, 1242–1250 (2014).

[69] Atashgahi, S., Sánchez-Andrea, I., Heipieper, H. J., van der Meer, J. R., Stams, A. J. M. and Smidt, H. Prospects for harnessing biocide resistance for bioremediation and detoxification. Science (80-.). 360, 743–746 (2018).

