## Supplementary Material for "*Escherichia coli* adapts metabolically to 6- and 7-fluoroindole, enabling proteome-wide fluorotryptophan substitution"

**Table of Contents of Supporting Information**

**Figures S1 – S12:**

**Tables S1 – S6:**

### 1. Analysis of 6- and 7-fluoroindole by GC-MS

Commercial preparations of 6- and 7-fluoroindole were purchased from Sigma Aldrich and specified with 99.9 % purity (6Fi via infrared spectrum) as well as 97.3 % purity (7Fi via GC). In order to exclude possible contamination with traces of indole, both compounds were additionally analyzed by gas chromatography coupled to mass spectrometry (GC-MS) in a targeted manner. Equal mixtures of indole and either 6-fluoroindole or 7-fluoroindole were used to develop a method with sufficient separation capacity (retention times are: indole 9.125 min, 6Fi 9.303 min and 7Fi 8.670 min). Furthermore, a calibration curve of indole was used to determine the limit of detection (LOD) of the method, which was found to be < 100 pmol. The GC-MS analysis was performed on an Agilent 5977 MSD system with the following parameters: column 5 % phenyl-methylpolysiloxane (Agilent 19091S-433UI), 325 °C, 30 m x 250 µm x 0.25 µm; gradient 50 °C (3 min isothermal), ramp 20 °C/min to 300 °C for 2 min; full-scan (50 – 350 m/z); injector 1 µL with 10:1 split ratio; flow rate 1 mL/min; pressure 7.6522 psi; heater 300 °C; EI mode; MS source: 230 °C, MS quad 150 °C. All measurements were performed at least twice.

Indole was used as positive control and for determination of the detection limit; it was characterized with a retention time of 9.125 min and m/z 117.1 as well as m/z 90.1 (**Figure S1**). Total ion current (TIC) chromatograms of 6- and 7-fluoroindole (**Figure S2, Figure S3**) were screened for contaminations, especially for adaptation disturbing non-fluorinated indole. Both TICs exhibit only one peak that could be assigned to the respective fluorinated indole, based on the mass spectrometric signals identified to be representative for the target substances (using NIST MS search 2.2 program). No indole could be detected within the LOD, hence the preparations of 6- and 7-fluoroindole were considered suitable for ALE. Furthermore, attempts to improve the analysis by *N,O*-Bis-(trimethylsilyl)-trifluoracetamid (BSTFA)-derivatization failed.

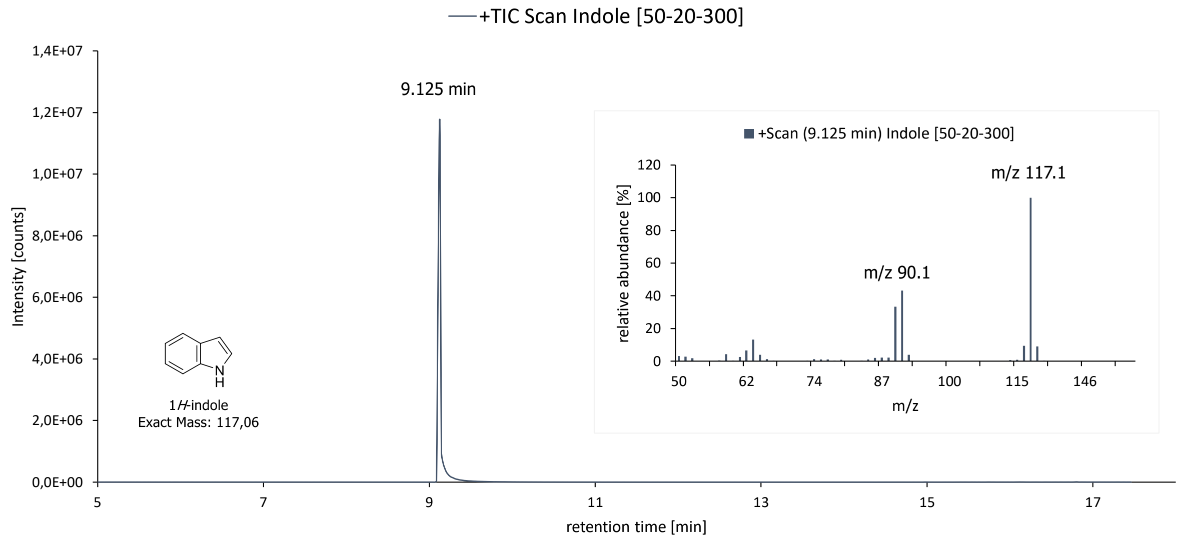

Figure S1. GC-MS full scan of indole. TIC chromatogram and ions detected at retention time 9.125 min; m/z 117.1 (100 %) and m/z 90.1 (43 %).

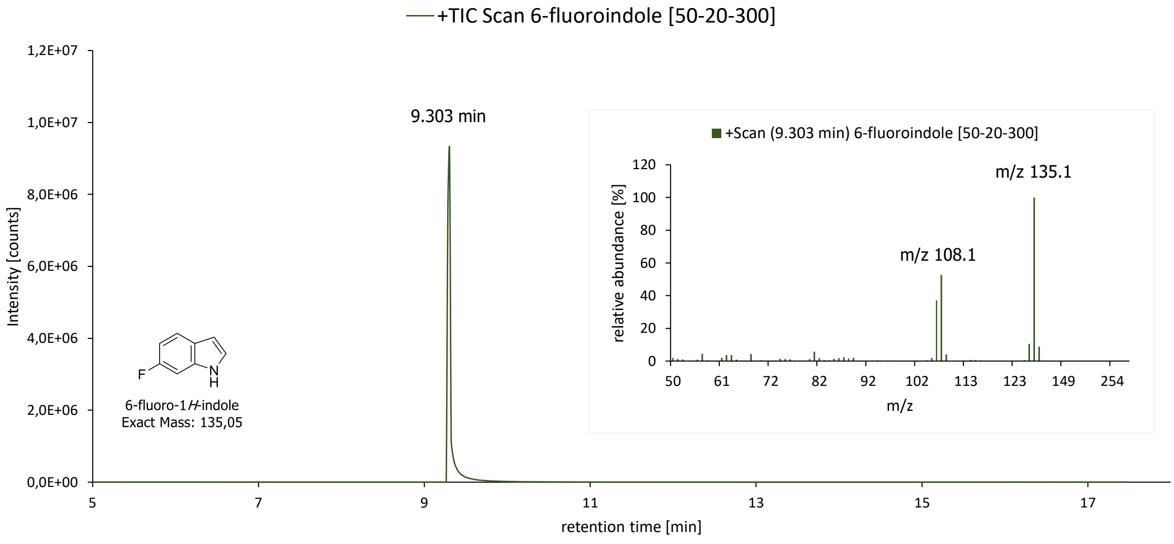

Figure S2. GC-MS full scan of 6-fluoroindole. TIC chromatogram and ions detected at retentions time 9.303 min; m/z 135.1 (100 %) and m/z 108.1 (53 %).

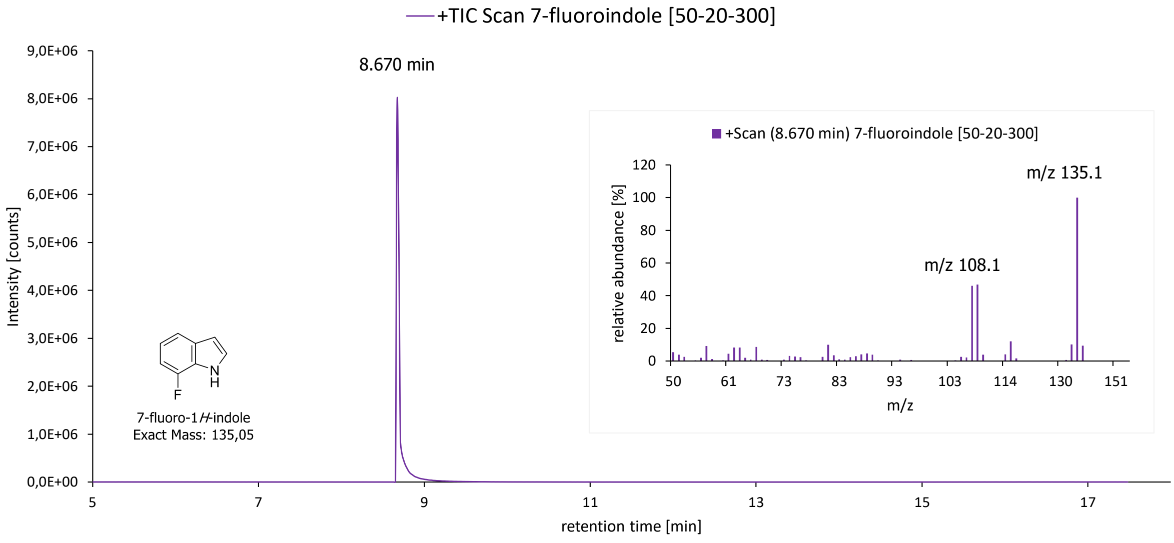

Figure S3. GC-MS full scan of 7-fluoroindole. TIC chromatogram and detected ions at retention time 8.670 min; m/z 135.1 (100 %) and m/z 108.1 (47 %).

### 2. Genetic configurations

Originating from *E. coli* MG1655, a strain with clean genomic background was designed by knocking out parts of the tryptophan operon (Δ*trpLEDC*) and the tryptophanase gene (Δ*tnaA*).[1] Before starting the evolution experiment, the genotypes of both strains were verified by standard PCR (Q5 Polymerase, New England Biolabs) and DNA sequencing (Microsynth AG, Switzerland). The analyses confirm the presence of *trpBA* encoding the TrpS enzyme in both strains as well as the deletion of the genes *trpLEDC* and *tnaA* in TUB00. According to the primer binding (**Figure S4**) for TUB00 shorter amplicons were detected for C4_trpB|C2_trpL (deletion *trpLEDC*) and C1_trpA|C2_trpL (full length amplification) and no band for C5_trpE|C2_trpE (no primer binding due to deleted *trpE*), as well as a shorter DNA fragment for full length amplification of the *tnaA* gene (C1_tnaA|C2_tnaA) and no band for C1_tnaA|C5_tnaA (no primer binding C5_tnaA due to deletion). This obligatory genetic constitution was also checked regularly during the entire adaptation process.

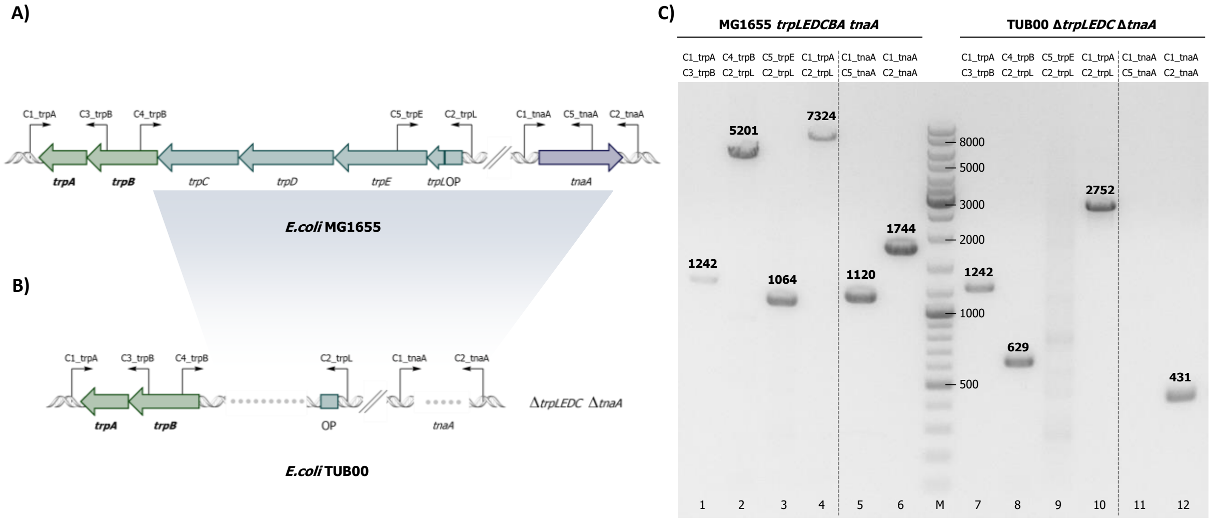

Figure S4. Overview of the genetic configuration. The genotypes of A) *E. coli* MG1655 and B) *E. coli* TUB00, as well as C) typical PCR pattern of both strains. A) and B) show the primer binding in the target genomic regions of the parent strain MG1655 with intact trp operon and tryptophanase and of the trp deficient derivative TUB00 with genetic deletions Δ*trpLEDC* Δ*tnaA*.

### 3. In vitro synthesis of 6- and 7-fluorotryptophan by the tryptophan synthase (TrpS)

A key aspect of our described ALE setup is the *in-situ* synthesis of fluorinated tryptophan from fluorinated indole. This corresponds to the last step of the endogenous Trp biosynthesis and is catalyzed by the tryptophan synthase (TrpS, *trpBA*); the reaction is a cofactor (pyridoxal 5'-phosphate, PLP) mediated condensation of L-serine and indole to yield Trp. Owing to its substrate promiscuity, TrpS also permits the conversion of fluorinated indole to fluorinated Trp. This reaction is described in literature for a variety of indole analogs, but we confirmed that also experimentally for our target substrates 6Fi and 7Fi, as well as Ind that serves positive control.

We used the TrpS enzyme of *Salmonella typhimurium*, whose beta-subunit differs in only 3.5 % from the *Escherichia coli* TrpS.[2] The reaction conditions were adapted from Wilcox protocol[3] . The optimized reaction (1 mL) encompassed 1 mM L-serine, 1 mM Indole or else 6- or 7-fluorondole, 0.8 mM pyridoxal-5'-phosphase (PLP) and 0.5 mM Trp synthase in 50 mM potassium phosphate buffer (pH 7.8). The mixture was incubated 24 h at 37 °C and the reaction progress was monitored by thin-layer chromatography (TLC) using ninhydrin staining to detect the amine moiety of the resulting amino acid; stationary phase: TLC silica gel 60 $F_{254}$, mobile phase: nBuOH : $CH_3COOH$ : $H_2O$ (2 : 1 : 1), UV = 254 nm. After purification via ion-exchange chromatography (IEX; Dowex 50WX8 50-100 mesh cationic resin, elution by ≈12 % $NH_3$), the conversion of the desired Trp or else 6FTrp or 7FTrp was confirmed by LC-ESI-MS (**Figure S5**). As controls, commercially available compounds were used (purchased from abcr). The calculated masses match to the observed and both signals (from *in vitro* and commercial samples) overlap.

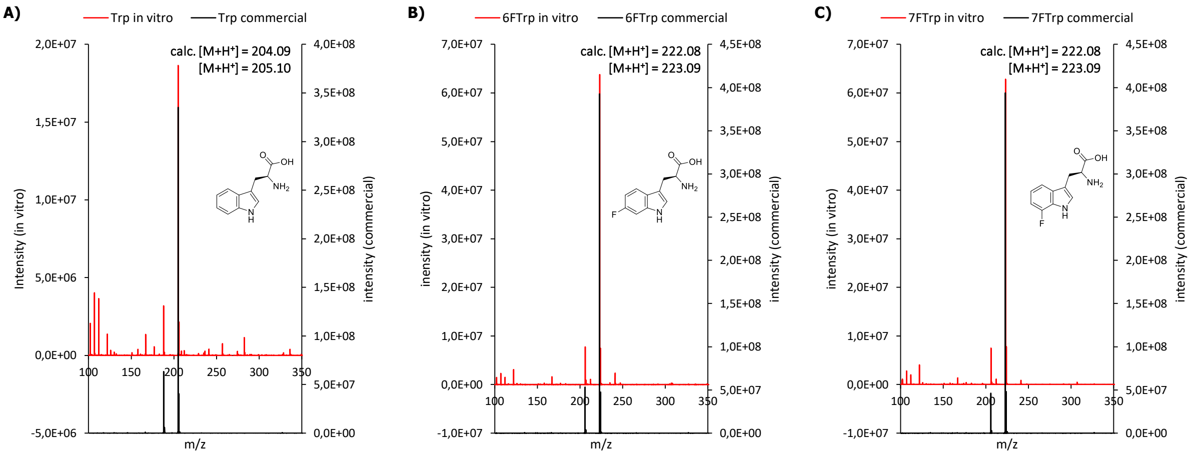

Figure S5. In vitro synthesis of Trp and FTrp. Mass spectrometric analyses (LC-ESI-MS) of A) Trp, B) 6FTrp and C) 7FTrp synthesized by TrpSMass spectra of the *in vitro* produced tryptophans are colored in red and those of the commercial tryptophans are depicted in black at the second y-axis.

### 4. Adaptive laboratory evolution (ALE) experiments

For the adaptation of *Escherichia coli* towards usage of 6- and 7-fluoroindole a serial transfer regime using batch cultures under aerobic conditions was applied. The cells were propagated in synthetic minimal medium (New Minimal Medium, NMM)[4] which consisted of: 7.5 mM $(NH_4)_2SO_4$, 8.5 mM NaCl, 22.5 mM $KH_2PO_4$, 50 mM $K_2HPO_4$, 1 mM $MgSO_4$, 20 mM D-glucose, 1 mg/L $Ca^{2+}$ (as $CaCl_2$), 1 mg/L $Fe^{2+}$ (as $FeCl_2$), 10 mg/L biotin, 10 mg/mL thiamine and 10 ng/L trace elements ($Cu^{2+}$, $Zn^{2+}$, $Mn^{2+}$, $MoO_4^{2-}$). In addition, the medium was variably enriched with indole (2.5 µM – 0 µM), fluoroindole (70 µM – 30 µM) and amino acids (50 mg/L), and was dynamically adjusted during the adaptation process (**Table S1-S4**).

The cultivation of each experiment (adaptation to 6Fi, 7Fi, W, Ind) was executed in parallel using three biological replicates (assumed to be isogenic) of our metabolic prototype TUB00 (Δ*trpLEDC* Δ*tnaA*), which is a derivative of *Escherichia coli* MG1655. 10 mL culture were incubated in 100 mL-shaking flasks at 30 °C, 180 rpm. When the cells reached their early stationary phase (usually after 50 - 65 h) fresh medium was inoculated to an $OD_{600}$ of 0.02 with the parent culture. The adjustment of the medium composition (increasing the selective pressure by depleting the indole and cAA concentration) was carried out when the cells proliferate sufficiently at least for two consequential passages under the parent conditions or no further growth improvement could be detected. The transfer steps of this serial dilution approach are indicated as “passages” and samples of each passage were preserved as cryo-stocks (25 % glycerol, -80 °C) for further analysis.

Beforehand ancestral TUB00 cultures were supplemented with different concentrations of indole (0 µM – 30 µM) and fluoroindole (5 µM – 1000 µM) to determine the starting conditions. 2.5 µM indole and 70 µM of 6Fi or 7Fi were found to allow the cells to grow to a sufficient optical density of $OD_{600}$ 0.7 – 1.0. Within the ALE setup, the concentration of fluoroindole was kept high; then first the concentration of indole was gradually reduced until complete depletion (ALE phase 1) and subsequently the amino acid supply was stepwise ceased (ALE phase 2). For the amino acid removal two different approached OC (overall concentration) and MB (metabolic blocks) were applied. At this stage the evolving lineages separated into either three (6TUB128-OC, 6TUB165-MB4, 6TUB165-MB3) or else two (7TUB165-OC, 7TUB165-MB) independent strains. In case of the 7-fluoroindole ALEs the growth conditions had to be relaxed (reduction of 7Fi concentration and increasing of cAA supply) in order to retain sufficient cell growth.

Table S1. New Minimal Medium (NMM) composition during the adaptation of 6TUB128-OC. NMM is designated as follows: NMM(a-b-c) where "a" corresponds to the amino acid supply, "b" to the 6-fluoroindole concentration [µM] and "c" to the indole concentration [µM]. Canonical amino acids (cAAs), except for Trp, were supplied as mixture (cAA19 mix, containing Tyr, Phe, Cys, Gly, Ser, Leu, Ile, Val, Ala, Asp, Asn, Glu, Gln, His, Met, Pro, Arg, Lys, Thr) in an initial concentration of 50 mg/L, which was gradually reduced in ALE phase 2 as depicted. Below 50 mg/L the cAA concentration is given in square brackets. The passages cultivated in the respective medium composition are provided.

| Medium (NMM) | cAA19 mixture [mg/L] | Indole [µM] | 6-fluoroindole [µ] | Passages |
|---|---|---|---|---|
| **19-70-2.5** | 50 | 2.5 | 70 | 1-3 |
| **19-70-1.0** | 50 | 1.0 | 70 | 4-5 |
| **19-70-0.5** | 50 | 0.5 | 70 | 6-18 |
| **19-70-0.25** | 50 | 0.25 | 70 | 19-25 |
| **19-70-0.10** | 50 | 0.10 | 70 | 26-35 |
| **19-70-0.05** | 50 | 0.05 | 70 | 36-40 |
| **19-70-0.01** | 50 | 0.01 | 70 | 41-44 |
| **19-70-0** | 50 | 0 | 70 | 45-53 |
| **[40]-19-70-0** | 40 | 0 | 70 | 54-58 |
| **[30]-19-70-0** | 30 | 0 | 70 | 59-62 |
| **[20]-19-70-0** | 20 | 0 | 70 | 63-66 |
| **[10]-19-70-0** | 10 | 0 | 70 | 67-71 |
| **[7.5]-19-70-0** | 7.5 | 0 | 70 | 72-77 |
| **[5.0]-19-70-0** | 5.0 | 0 | 70 | 78-81 |
| **[2.5]-19-70-0** | 2.5 | 0 | 70 | 82-84 |
| **0-70-0** | 0 | 0 | 70 | 85-128 |

16 adaptation steps

Table S2. New Minimal Medium (NMM) composition during the adaptation of 6TUB165-MB4 and 6TUB165-MB3. NMM is designated as follows: NMM(a-b-c) where "a" corresponds to the amino acid supply, "b" to the 6-fluoroindole concentration [µM] and "c" to the indole concentration [µM]. Canonical amino acids (cAAs; F, Y, A, V, L, I, H, G, S, C, K, D, N, M, T, P, E, Q, R), except for Trp, were supplied in separate stock solutions (50 mg/L) and removed as metabolic blocks in ALE phase 2. For 6TUB165-MB4 and 6TUB165-MB3 different routes for the cAA elimination were used as depicted. The passages cultivated in the respective medium composition are provided.

| Medium (NMM) | cAAs supplied | Indole [µM] | 6-fluoroindole [µ] | Passages |
|---|---|---|---|---|
| **19-70-2.5** | FY AVLI H GSC KDNMT PEQR | 2.5 | 70 | 1-3 |
| **19-70-1.0** | FY AVLI H GSC KDNMT PEQR | 1.0 | 70 | 4-5 |
| **19-70-0.5** | FY AVLI H GSC KDNMT PEQR | 0.5 | 70 | 6-18 |
| **19-70-0.25** | FY AVLI H GSC KDNMT PEQR | 0.25 | 70 | 19-25 |
| **19-70-0.10** | FY AVLI H GSC KDNMT PEQR | 0.10 | 70 | 26-35 |
| **19-70-0.05** | FY AVLI H GSC KDNMT PEQR | 0.05 | 70 | 36-40 |
| **19-70-0.01** | FY AVLI H GSC KDNMT PEQR | 0.01 | 70 | 41-44 |
| **19-70-0** | FY AVLI H GSC KDNMT PEQR | 0 | 70 | 45-53 |
| **17-70-0** | AVLI H GSC KDNMT PEQR | 0 | 70 | 54-59 |
| **13-70-0** | H GSC KDNMT PEQR | 0 | 70 | 60-63 |
| **12-70-0** | GSC KDNMT PEQR | 0 | 70 | 64-67 |
| **9-70-0** | KDNMT PEQR | 0 | 70 | 68-72 |
| **6TUB165-MB4** | | | | |
| **6-70-0** | KD PEQR | 0 | 70 | 73-77 |
| **4-70-0** | PEQR | 0 | 70 | 78-81 |
| **0-70-0** | 0 | 0 | 70 | 82-165 |
| **6TUB165-MB3** | | | | |
| **5-70-0** | KDMNT | 0 | 70 | 73-77 |
| **3-70-0** | NMT | 0 | 70 | 78-81 |
| **0-70-0** | 0 | 0 | 70 | 82-165 |

15 adaptations steps

Table S3. New Minimal Medium (NMM) composition during the adaptation of 7TUB165-OC. NMM is designated as follows: NMM(a-b-c) where "a" corresponds to the amino acid supply, "b" to the 7-fluoroindole concentration [µM] and "c" to the indole concentration [µM]. Canonical amino acids (cAAs), except for Trp, were supplied as mixture (cAA19 mix, containing Tyr, Phe, Cys, Gly, Ser, Leu, Ile, Val, Ala, Asp, Asn, Glu, Gln, His, Met, Pro, Arg, Lys, Thr) in an initial concentration of 50 mg/L, which was gradually reduced in ALE phase 2 as depicted. Below 50 mg/L the cAA concentration is given in square brackets. The passages cultivated in the respective medium composition are provided. During this adaptation the medium composition need several times relaxed (reduction of 7Fi concentration, increase of cAA supply) to uphold cell growth.

| Medium (NMM) | cAA19 mixture [mg/L] | Indole [µM] | 7-fluoroindole [µM] | Passages |
|---|---|---|---|---|
| **19-70-2.5** | 50 | 2.5 | 70 | 1-2 |
| **19-70-1.0** | 50 | 1.0 | 70 | 3-6 |
| **19-70-0.5** | 50 | 0.5 | 70 | 7-14 |
| **19-70-0.25** | 50 | 0.25 | 70 | 15-18 |
| **19-70-0.10** | 50 | 0.10 | 70 | 19-26 |
| **19-70-0.05** | 50 | 0.05 | 70 | 27-35 |
| **19-70-0.01** | 50 | 0.01 | 70 | 36-41 |
| **19-70-0** | 50 | 0 | 70 | 42-53 |
| **[40]-19-70-0** | 40 | 0 | 70 | 54-58 |
| **[30]-19-70-0** | 30 | 0 | 70 | 59-62 |
| **[20]-19-70-0** | 20 | 0 | 70 | 63-66 |
| **[10]-19-70-0** | 10 | 0 | 70 | 67-71 |
| **[7.5]-19-70-0** | 7.5 | 0 | 70 | 72-80 |
| **[5.0]-19-70-0** | 5.0 | 0 | 70 | 81-83 |
| **[10]-19-50-0** | 10 | 0 | 50 | 84-88 |
| **[5.0]-19-50-0** | 5.0 | 0 | 50 | 89-92 |
| **[2.5]-19-50-0** | 2.5 | 0 | 50 | 93-100 |
| **[2.5]-19-30-0** | 2.5 | 0 | 30 | 101-102 |
| **[10]-19-30-0** | 10 | 0 | 30 | 103-104 |
| **[5.0]-19-30-0** | 5.0 | 0 | 30 | 105-106 |
| **[2.5]-19-30-0** | 2.5 | 0 | 30 | 107-109 |
| **[1.0]-19-30-0** | 1.0 | 0 | 30 | 110-111 |
| **[1.5]-19-30-0** | 1.5 | 0 | 30 | 112-125 |
| **[1.0]-19-30-0** | 1.0 | 0 | 30 | 126-128 |
| **[0.5]-19-30-0** | 0.5 | 0 | 30 | 129-131 |
| **0-30-0** | 0 | 0 | 30 | 132-143 |
| **0-70-0** | 0 | 0 | 70 | 144-165 |

27 adaptation steps

Table S4. New Minimal Medium (NMM) composition during the adaptation of 7TUB165-MB. NMM is designated as follows: NMM(a-b-c) where "a" corresponds to the amino acid supply, "b" to the 7-fluoroindole concentration [µM] and "c" to the indole concentration [µM]. Canonical amino acids (cAAs; F, Y, A, V, L, I, H, G, S, C, K, D, N, M, T, P, E, Q, R), except for Trp, were supplied in separate stock solutions (50 mg/L) and removed as metabolic blocks or else one by one in ALE phase 2. The passages cultivated in the respective medium composition are provided. The medium composition of this adaption also had to be relaxed multiple times by decreasing the 7Fi concentration and increasing the number of amino acids supplied.

| Medium (NMM) | cAAs supplied | Indole [µM] | 6-fluoroindole [µ] | Passages |
|---|---|---|---|---|
| **19-70-2.5** | FY AVLI H GSC KDNMT PEQR | 2.5 | 70 | 1-2 |
| **19-70-1.0** | FY AVLI H GSC KDNMT PEQR | 1.0 | 70 | 3-6 |
| **19-70-0.5** | FY AVLI H GSC KDNMT PEQR | 0.5 | 70 | 7-14 |
| **19-70-0.25** | FY AVLI H GSC KDNMT PEQR | 0.25 | 70 | 15-18 |
| **19-70-0.10** | FY AVLI H GSC KDNMT PEQR | 0.10 | 70 | 19-26 |
| **19-70-0.05** | FY AVLI H GSC KDNMT PEQR | 0.05 | 70 | 27-35 |
| **19-70-0.01** | FY AVLI H GSC KDNMT PEQR | 0.01 | 70 | 36-41 |
| **19-70-0** | FY AVLI H GSC KDNMT PEQR | 0 | 70 | 42-53 |
| **17-70-0** | AVLI H GSC KDNMT PEQR | 0 | 70 | 54-58 |
| **12-70-0** | GSC KDNMT PEQR | 0 | 70 | 59-83 |
| **12-50-0** | GSC KDNMT PEQR | 0 | 50 | 84-92 |
| **11-50-0** | GSC DNMT PEQR | 0 | 50 | 93-97 |
| **10-50-0** | GSC NMT PEQR | 0 | 50 | 98-100 |
| **10-30-0** | GSC NMT PEQR | 0 | 30 | 101-102 |
| **13-30-0** | H GSC KDNMT PEQR | 0 | 30 | 103-104 |
| **12-30-0** | GSC KDNMT PEQR | 0 | 30 | 105-106 |
| **11-30-0** | GSC DNMT PEQR | 0 | 30 | 107-108 |
| **10-30-0** | GSC DMT PEQR | 0 | 30 | 109-113 |
| **9-30-0** | GSC DM PEQR | 0 | 30 | 114-115 |
| **8-30-0** | GSC M PEQR | 0 | 30 | 116-117 |
| **7-30-0** | SC M PEQR | 0 | 30 | 118-120 |
| **6-30-0** | S M PEQR | 0 | 30 | 121-125 |
| **5-30-0** | M PEQR | 0 | 30 | 126-128 |
| **4-30-0** | M PEQ | 0 | 30 | 129-131 |
| **3-30-0** | M EQ | 0 | 30 | 132-133 |
| **2-30-0** | M E | 0 | 30 | 134-135 |
| **1-30-0** | M | 0 | 30 | 136-150 |
| **0-30-0** | 0 | 0 | 30 | 151-154 |
| **0-70-0** | 0 | 0 | 70 | 155-165 |

30 adaptation steps

### 5. Adaptation towards 6Fi and 7Fi

Description of the 6Fi adaptation course: ALE phase 1, for 6TUBX-OC and the two 6TUBX-MB lineages comprised 53 passages that means 301 generations in 128.8 days. We propagated the 6TUBX-OC lineage for a total of 128 passages (780 generations, 317.2 days). These cells completed phase 2 after 31 passages (205 generations, 77.2 days) and grew 44 passages (274 generations, 111.2 days) in the final adaptation medium (NMM0-70-0). In contrast, the two 6TUBX-MB lineages (6TUB165-MB4 and 6TUB165-MB3) grew significantly longer with 165 passages (1024 generations, 410.1 days). They ended phase 2 after 28 passages (179 generations, 71.7 days) and grew twice as long in the final adaptation medium (NMM0-70-0) with 84 passages (544 generations, 209.6 days).

Description of the 7Fi adaptation course: The mutual phase 1 lasted 53 passages (301 generations, 122.1 days). The 7TUB165-OC lineage grew 79 passages (330 generations, 198.4 days) in phase 2, 33 passages (147 generations, 82.5 days) in phase 3 (12 passages (47 generations, 29.9 days) thereof with reduced 7Fi concentration in NMM0-30-0) and took 165 passages (778 generations, 403.0 days) for the entire adaptation process. A very similar trajectory was observed for 7TUB165 MB, whose development required 98 passages (397 generations, 243.9 days) in phase 2, 14 passages (54 generations, 35.0 days) in phase 3 (whereby a transition from 4 passages comprising 11 generations within 10 days in NMM0 30-0 was necessary); and comprises again a total of 165 passages (752 generations, 401 days). Since there was no further significant growth improvement expected, the adaptation was terminated after only 14 and 33 passages in phase 3 (NMM0-70-0), respectively. An overview of the ALE parameters (number of passages, number of generations and incubation time in days) is given in **Figure S6** and **Table S5**.

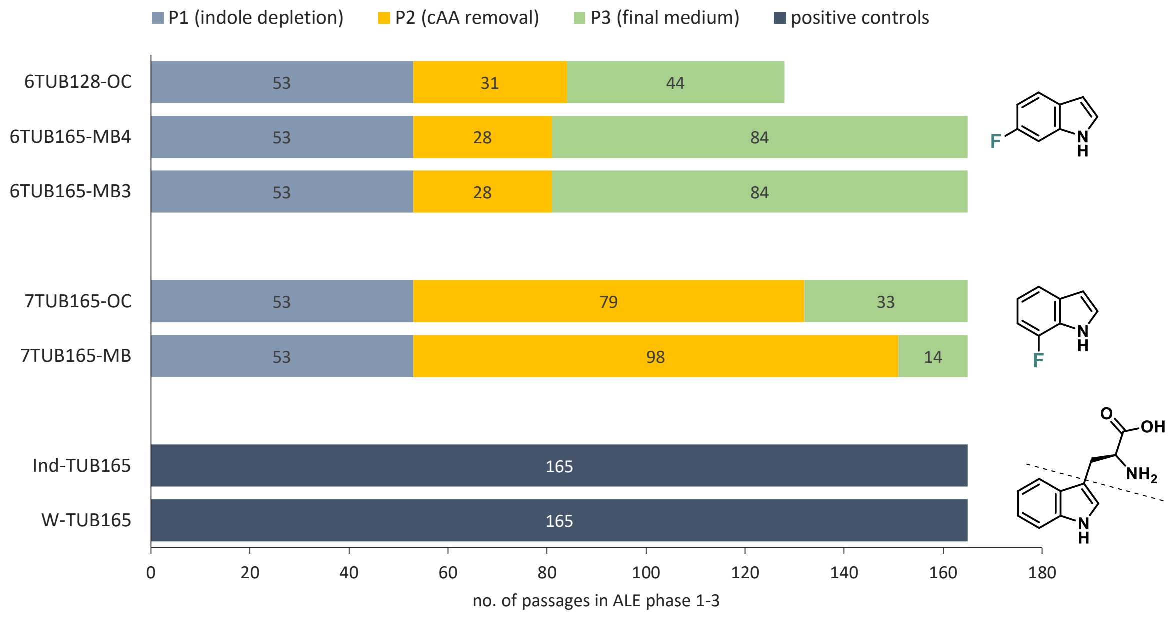

Figure S6. Overview of ALE trajectories. The extent of the ALE phases is illustrated by means of passages needed to complete each adaptation phase (1, 2) and the number of passages spent under final growth conditions (phase 3). The indole depletion (phase 1, grey) took 53 passages for each ALE. The cAA removal (phase 2, yellow) was accomplished in about 30 passages for the 6Fi ALEs as well as 79 and 98 passages, respectively, for the 7Fi ALEs. And the 6Fi adapted cells were grown for 44 or 84 passages in their final medium composition (phase 3, green), whereas the 7Fi adapted strains were hold only 14 or 33 passages under final adaptation conditions. The positive controls were not subjected to the ALE phase concept but cultivated continuously for equally 165 passages in NMM0-0-70 (Ind or Trp).

Table S5. Overview of the technical details of the adaption trajectories. Attributes passage number (pass.), number of generations (gen.) and cultivation time in days for every ALE phase (P1-3) and the entire adaption process are shown.

| | | ***Entire adaptation*** | | | ***P1 (indole depletion)*** | | | ***P2 (cAA removal)*** | | | ***P3 final medium*** | | |
|---|---|---|---|---|---|---|---|---|---|---|---|---|---|
| | | pass. | gen. | days | pass. | Gen. | days | pass. | gen. | days | pass. | gen. | days |
| ***W-TUB165*** | W | 165 | 1273 | 413.8 | | | | | | | | | |
| ***Ind-TUB165*** | Ind | 165 | 1270 | 413.6 | | | | | | | | | |
| ***6TUB128-OC*** | 6Fi | 128 | 780 | 317.2 | 53 | 301 | 128.8 | 31 | 205 | 77.2 | 44 | 274 | 111.2 |
| ***6TUB165-MB4*** | 6Fi | 165 | 1024 | 410.1 | 53 | 301 | 128.8 | 28 | 179 | 71.7 | 84 | 544 | 209.6 |
| ***6TUB165-MB3*** | 6Fi | 165 | 1024 | 410.1 | 53 | 301 | 128.8 | 28 | 179 | 71.7 | 84 | 544 | 209.6 |
| ***7TUB165-OC*** | 7Fi | 165 | 778 | 403.0 | 53 | 301 | 122.1 | 79 | 330 | 198.4 | 33 | 147 | 82.5 |
| ***7TUB165-MB*** | 7Fi | 165 | 752 | 401.0 | 53 | 301 | 122.1 | 98 | 397 | 243.9 | 14 | 54 | 35.0 |

While the 6Fi adapted cells reached growth in their final medium composition (NMM0-70-0) after around 80 passages, those adapted to 7Fi took with either 130 passages (7TUB165-OC) or 150 passages (7TUB165-MB) much longer to reach this stage. But we used this opportunity to prolong the propagation of the 6Fi lineages in ALE phase 3 to evaluate if the growth behavior would improve or an indole rejecting phenotype could evolved after all; i.e., 44 passages for 6TUB128-OC and 84 passages for 6TUB165-MB4 and -MB3 in NMM0-70-0, respectively. However, neither a significant improvement of the growth performance nor an unreserved fluoroindole preferring variant could be observed/isolated. Finally, also both 7Fi adapted strains grew stable in NMM0-70-0 for 10 passages; and since the population size did not further increase, we decided to terminate all adaptation experiments at this point. This ensured the reproducibility of the cell growth in medium supplemented only with fluoroindole but in absence of canonical indole and amino acids.

### 6. Adaptation of positive controls

The same procedure as for the Fi-adaptation was followed with the positive controls, adapted to grow on 70 µM of either indole (Ind-TUBX) or tryptophan (W-TUBX). These cells were cultivated under the same growth regime for 165 passages, except for the amino acid removal process of ALE phase 2 (NMM19-0-70), which was omitted (**Figure S7**).

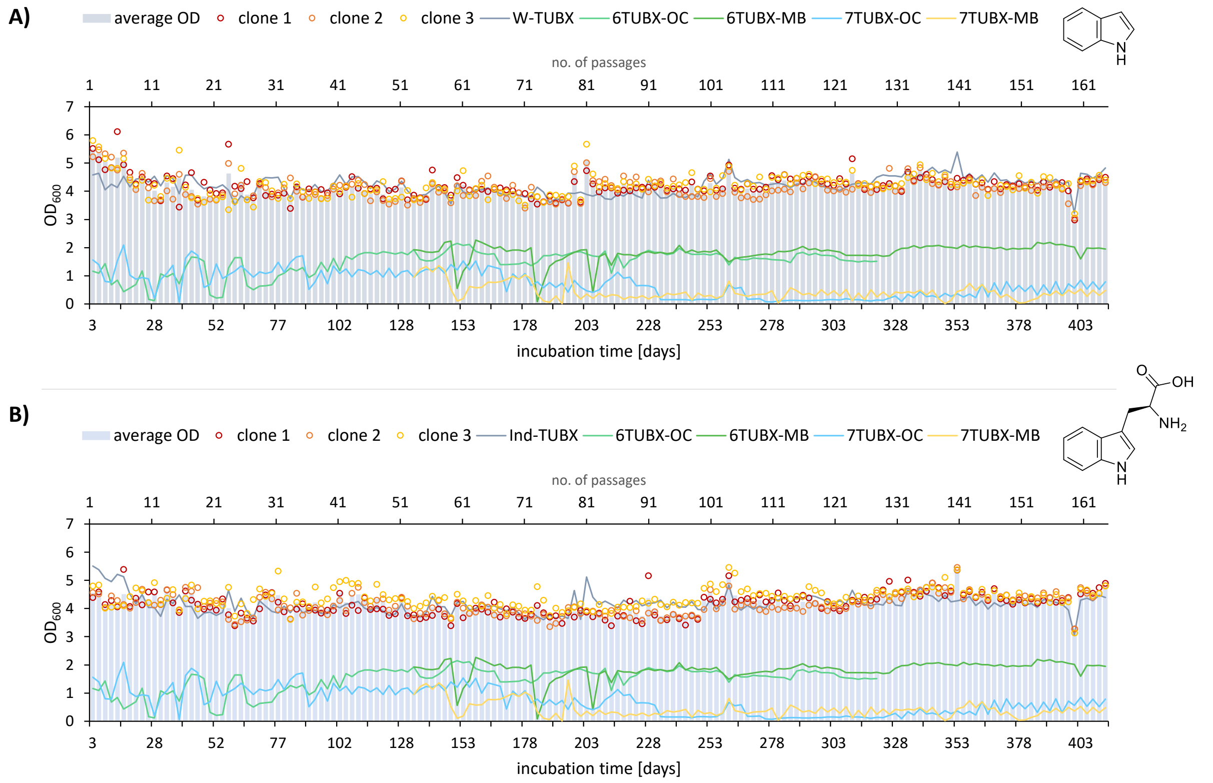

Figure S7. Cultivation schemes of the positive controls. Adaptation of *E. coli* towards A) indole and B) tryptophan is shown. The optical density ($OD_{600}$) is plotted against the incubation time and the number of passages (reinoculations steps); the average $OD_{600}$ is shown as bar plot and the individual clones are represented as circles. The average OD of corresponding positive controls (W-TUBX or Ind-TUBX in grey) as well as 6TUBX-OC (turquoise) 6TUBX-MB (green) and 7TUBX-OC (blue), 7TUBX-MB (yellow) lineages are depicted as lines. The cells grew constantly in NMM19-0-70 supplemented with 19 cAAs (50 mg/L) except for Trp and either 70 µM Trp or Ind as target substrate; NMM is designated as follows: NMM(a-b-c) where "a" is the supplied amino acids, "b" the concentration of fluoroindole in µM and "c" the concentration of indole in µM.

### 7. Global Trp for F-Trp substitution (proteomics)

Final isolates of the adapted strains 6TUB128-OC, 6TUB165-MB4, 6TUB165-MB3, 7TUB165-OC, 7TUB165-MB and positive controls Ind-TUB165, W-TUB165 as well as TUB00 were grown in the respective NMM supplemented with either 70 µM indole, 6Fi or 7Fi. Exponential growing cells were harvested by centrifugation (18000 g, 4 °C, 20 min) and pellets were stored at -20 °C. *E. coli* extracts were briefly run into an SDS-gel until the dye front had migrated about 5 mm into the gel and only the 3 mm behind the dye front were used for protein digestions. This step removes all positively charged molecules, small highly-mobile negatively charged ions, uncharged molecules and very large agglomerates that did not enter the gel. Following a protocol described by Kublik *et. al.*,[5] the gel slices were washed with $ddH_2O$, and the proteins were reduced with dithiothreitol to break disulfide bridges. To protect the reduced cysteine residues, carbamidylation using iodoacetamide was performed. Subsequently, an overnight trypsin digestion was initiated to generate peptides. The resulting peptides were extracted from the gel pieces and desalted using C18-tips through zip-tipping. These extracted peptide samples were then analyzed by nLC-MS/MS on a nanoUPLC system (nanoAcquity, Waters) coupled to an Orbitrap Fusion mass spectrometer (Thermo Scientific), as previously described.[6] Protein identification was achieved using Proteome Discoverer (v2.4, Thermo Fisher Scientific) with the *E. coli* fasta genome database and SequestHT as the search engine, setting a false discovery rate threshold of 1 % for peptide identification using the Target Decoy PSM Validator node. The abundance of proteins was quantified through label-free quantification based on intensity values in precursor scans using the Minora node in Proteome Discoverer. For the identification of peptides, carbamidomethylation of cysteine residues was taken as a fixed modification, oxidation of methionine residues and fluorination of tryptophane residues were taken as dynamic modifications. Fluorination was calculated as the substitution of a hydrogen atom by fluor giving a mass change of +17.991 (**Figure S8**).

Within the limits of the method, global substitution of Trp by F-Trp was unambiguous proven in the adapted strains. The following false positives rates (mean of three biological replicates) were determined TUB00 (0.0225 % ±0.0318), Ind-TUB165 (0.0878 % ±0.0098), W-TUB165 (0.0485 % ±0.0485), 6TUB128-OC (0.2309 % ±0.1315), 6TUB165-MB4 (0.1702 % ±0.0555), 6TUB165-MB3 (0.1269 % ±0.1269), 7TUB165-OC (0.3984 % ±0.0822) and 7TUB165-MB (0.5154 % ±0.1173).

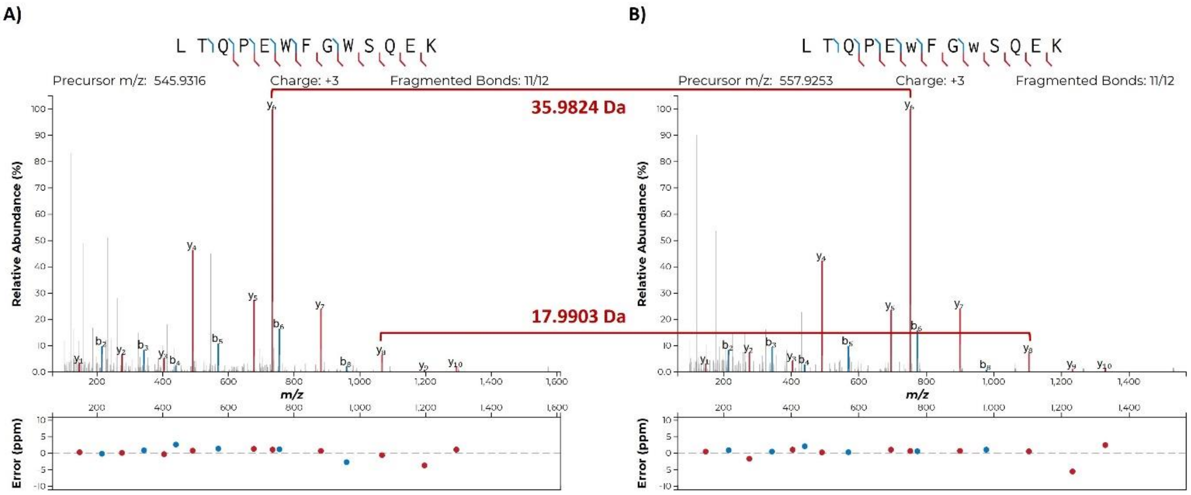

Figure S8. Example MS/MS spectra for the global Trp to FTrp substitution. The MS/MS spectra of oligopeptide ABC transporter periplasmic binding protein (NP_415759.1) peptide 352-LTQPEWFGWSQEK-366 from ancestral/positive controls (A) and FW-adapted strains (B) are shown, with and without fluorine modification on W6 and W9, respectively. The selected monoisotopic precursor m/z (z=+3) is given in each diagram and all detected y fragment ions are shown in red and b fragment ions are shown blue. Red brackets visualize the consistent m/z difference of 17.99 Da per fluorine substitution between equivalent y ions; $y_8$ (WSQEK) = 17.9903 Da for W8 fluorine modification and $y_6$ (WFGWSQEK) = 35.9824 Da for W6 and W9 fluorine modifications.

### 8. Subpopulation screening for an indole rejecting phenotype

Routine examination of the growth behavior of the 6Fi adapted strains in both indole and 6-fluorindole supplemented medium revealed that growth appeared to be reduced on indole compared to 6Fi. Upon this finding we designed a simple experiment to validate the growth behavior under different conditions, in terms of growth matrix (solid, liquid) and substrates (Ind, 6Fi). For this purpose, population samples of clone 1 of passage 124 of the three 6Fi-ALEs (6TUB124-OC, 6TUB124-MB4 and 6TUB124-MB3) were chosen and spread out for clone isolation. 20 single clones of every lineage were picked, diluted in 20 µL water and 1 µL was spotted on two agar plates containing either 6-fluorindole or indole (1:1 mixture of 2x $H_2O$ agar (9 %) and 2x NMM, either 19-0-70 (Ind) and 19-70-0 (6Fi)). We decided for relaxed growth conditions by means of adding cAAs to provoke any growth (desired as well as unwanted).

After incubation for two days at 30 °C, the agar plates showed that clones of the OC lineage grow well on both substrates, whereas the clones of the two MB lineages showed a preference for 6Fi. In this case there was no growth observed on the indole plate, which hints for the development of a changing nutritional preference in favor of the fluorinated substrates. Since growth conditions are in general more stringent in liquid culture than on plates, we repeated this setup in liquid media. Therefore, from the "19-70-0"-plate again 20 promising clones were selected out of the 60, washed twice in NMM0-0-0 to remove residual 6Fi and used to inoculate media (2 mL to $OD_{600}$ 0.02) again containing either 6Fi or indole (NMM19-0-70 (Ind) and NMM19-70-0 (6Fi)). The initial results were confirmed in liquid media, that the cells of the MB lineages acquired an increased preference for the fluorinated substrate as they reach higher cell densities (**Figure S9**). These experiments were repeated with cells of a subpopulation that were exposed to increased selection pressure; they are passages 140 (85 µM 6Fi) and 150 (100 µM 6Fi). Based on these experiments we stopped the adaptation of the OC-lineage as, in contrast to the MB lineages, the development of the desired phenotype no longer seemed to be expectable.

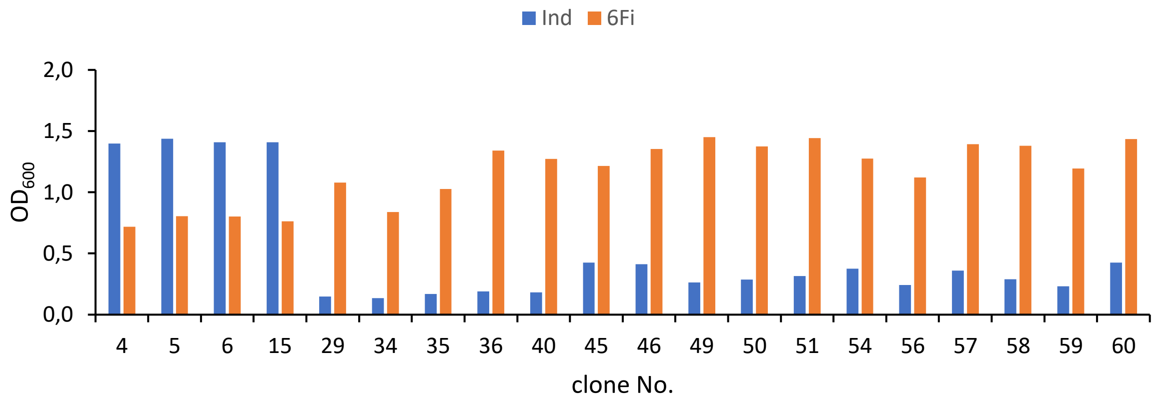

Figure S9. Subpopulation screening. Comparison of the growth behavior of selected clones in liquid media NMM19-0-70 (Ind) and NMM19 70-0 (6Fi). Clones 4, 5, 6 and 15 are derived from 6TUB124-OC, clones 29, 34, 35, 36 and 40 from 6TUB124-MB4 and clones 46, 49, 50, 51, 54, 56, 57, 59 and 60 are originated from 6TUB124-MB3. Cultures were incubated at 30 °C, 180 rpm for 24 h.

### 9. Growth curves

Isolates of the ancestral strain TUB00 and of the final isolates W-TUB165, Ind-TUB165, 6TUB128-OC, 6TUB165-MB4, 6TUB165-MB3, 7TUB165-OC and 7TUB165-MB were subjected to comprehensive growth analysis. For this purpose, growth curves in different medium compositions along the adaptation curse were recoded (**Figure S10**). These comprised LB (rich, undefined medium), NMM19-0-70 (Ind), NMM19-70-1 (6Fi or 7Fi, Ind), NMM19-70-0 (6Fi or 7Fi) and NMM0-70-0 (6Fi or 7Fi). Thereby, 6Fi adapted cells could be directly resuscitated in NMM0-70-0 (6Fi) from cryo stock, whereas 7Fi adapted cells had to be revived in nutrient rich LB medium, followed by two-times serial inoculation in their final adapted medium NMM0-70-0 (in order to restore the intracellular trp-free environment). Subsequently, the cells were washed twice with NMM0-0-0 to remove residual Ind, 6Fi or 7Fi, normed to $OD_{600}$ = 1, and then 200 µL of the respective medium composition were inoculated to $OD_{600}$ = 0.02. The growth was monitored in a 96-well plate (Greiner, flat bottom clear), sealed with breathable membrane (Breathe-Easy sealing membrane, Sigma Aldrich) at 30 °C with continuous shaking using a microplate reader (Tecan Infinite M200) that measured the absorbance at 600 nm in 10 min intervals. The measurements were performed at least twice and in triplicates (biological replicates).

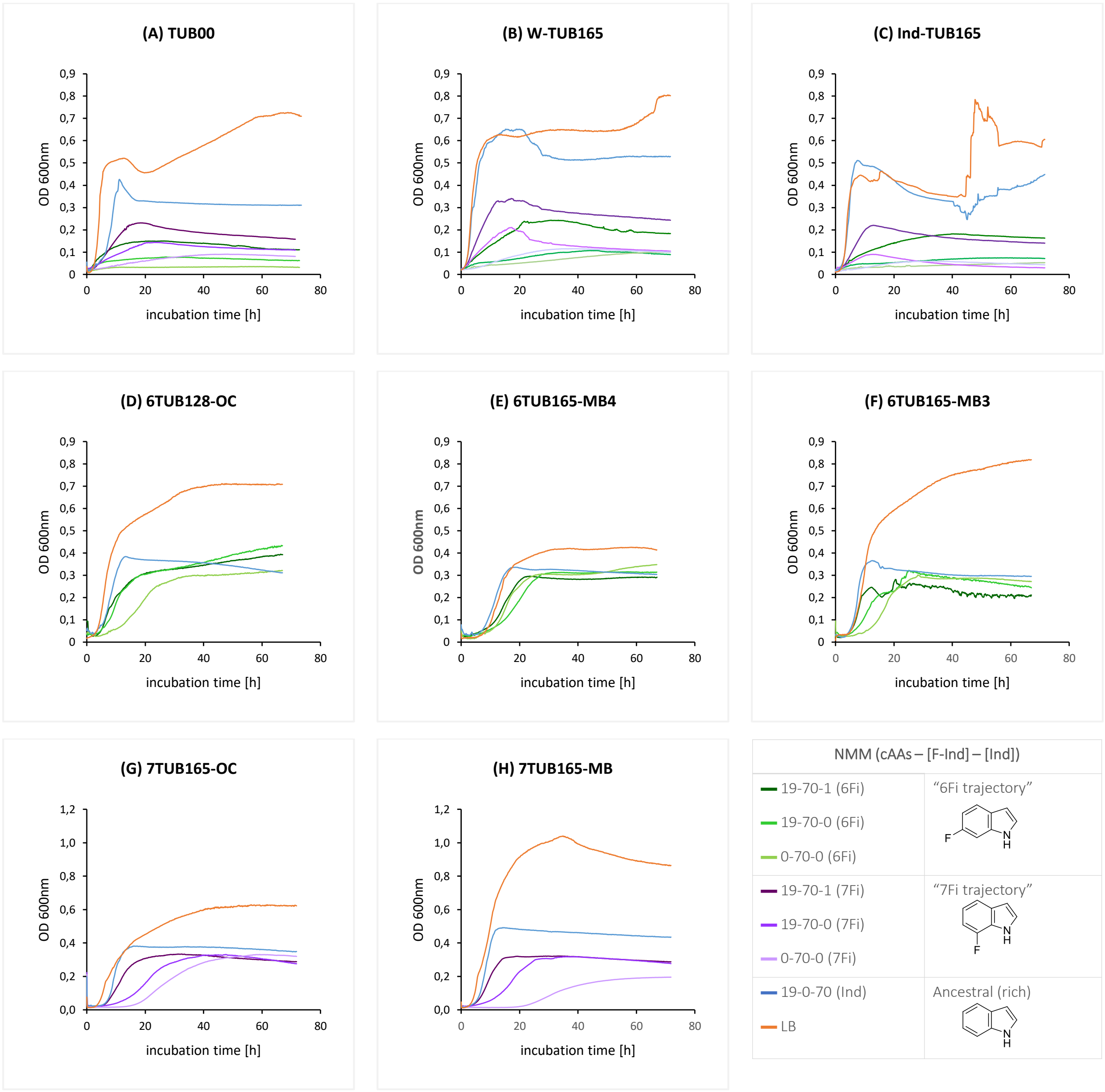

Figure S10. Growth curves of the ancestral strain and positive controls (A-C), the 6Fi adapted lineages (D-F) and the 7Fi adapted lineages (G-H), the final isolates respectively. The absorption at 600 nm is plotted against the incubation time [h] at 30 °C. Growth curves gained in LB are colored orange, in NMM19-0-70 (Ind) are blue, in 6Fi supplemented NMM are in green shades and in 7Fi supplemented NMM are colored in purple shades (see legend). The frizzy and instable curve parts obtained in C (LB and NMM19-0-70 (Ind)) and F (NMM19-70-1 (6Fi)) are rated as negligible, since for estimation of the growth behavior and calculation of the growth parameters only the exponential part of the curve is of interest.

As expected, the ancestral strain TUB00 (**Figure S10A**) reaches the highest density in rich medium, whereby the growth in common LB medium is better than in defined indole containing NMM (19-0-70). The growth in medium along the adaptation trajectory decreases in correlation with the nutrition supply, which was also expected by the wild type strain. In contrast, both positive controls W-TUB165 (**Figure S10B**) and Ind-TUB165 (**Figure S10C**) exhibit improved growth in NMM19-0-70 (Ind), which is comprehensible because these strains were adapted to grow on minimal medium. Interestingly, the trp and indole driven strains tolerate 7Fi better than 6Fi, although based on the adaptation results, we had expected the opposite result.

Among the 6Fi-adapted lineages 6TUB128-OC, 6TUB165-MB4 and 6TUB165-MB3 (**Figure S10D-F**) the growth with NMM19-0-70 (Ind) as well as along the adaptation trajectory (with 6Fi supplemented NMM) is relatively equal. However, the growth of the MB4 lineage is the most striking since here, all growth curves are very close to each other. In total the MB4 lineage seems to be best adapted to 6Fi, indicated by the highest growth rate in the final medium and by the comparatively low growth in rich LB medium, which in turn could indicate an approaching rejection of the indole and need for 6-fluoroindole.

For the 7Fi adapted cells, there are clear differences between the OC and MB lineages (**Figure S10G-H**). The OC lineage appears to be better adapted as the cells have a significantly shorter lag time and reach higher optical densities; a behavior that is especially pronounced in the final medium composition NMM0-70-0. This correlates also with the fact that this lineage was cultivated much more passages in the final medium compared to the MB-lineage, where the amino acid removal was more struggling. Overall, the response to media composed of 7Fi exhibits the same trend for both lineages; according to the nutrition score, the growth rate also decreases with reduced nutrient content (Ind, cAAs), but the maximum attainable optical density stays relatively constant. This behavior indicates poor adaptation, although this was already evident from the cultivation schemes. In any case, these cell lines prefer the ancestral medium over the fluorinated one and grow worst in their final medium.

### 10. Viability assay (CCK-8)

The cell viability was determined as additional parameter for bacterial fitness of the adapted strains; and was assessed using cell counting kit-8 (CCK-8; Sigma-Aldrich), which is based on dehydrogenase activity detection in viable cells. For the proliferation assay 190 µL of 5.0 x $10^{07}$ cells/mL suspension were seeded, added with 10 µL of the CCK-8 solution and incubated for 4 h at 30 °C without shaking. Then, the absorbance at 450 nm was measured using microplate reader (Tecan reader M200), which is directly proportional to the number of proliferating cells. As background control medium without cells was used and the viability of TUB00 was set 100 % in comparison to the adapted strains. Measurements were performed in triplicates and repeated at least twice.

### 11. Determination of the minimal inhibitory concentration of vancomycin

For testing the antibiotic susceptibility, the minimal inhibitory concentration (MIC) of vancomycin was determined (**Figure S11**). Isolates from cryo stocks of the respective strains were resuscitated either in 2-3 mL LB or appropriate NMM, incubated at 30 °C, 200 rpm and inoculated into NMM if necessary (see above section 9). Subsequently, cells were used to inoculate 3 mL NMM in culture tubes as follows NMM 0-0-70 (Ind or Trp) for TUB00, W-TUB165, Ind-TUB165 and NMM 0-70-0 (6Fi or 7Fi) for adapted strains 6TUB128-OC, 6TUB165-MB4, 6TUB165-MB3 and 7TUB165-OC, 7TUB165-MB. For determination of the MIC a cell density corresponding to 0.5 McFarland standard was used and cells were treated with serial dilutions of vancomycin (0, 25, 50, 100, 200, 400 µg/mL). Cell growth (incubation at 30 °C, 200 rpm) was monitored by measuring the $OD_{600}$ every 24 hours and MIC was defined as the lowest concentration at which no increase in cell density could be observed ($OD_{600}$ = 0.1). Measurements were performed using biological triplicates.

Generally, the cell density of all tested strains decreases with increasing vancomycin concentration, however the susceptibility is as follows: 6TUB128-OC, 6TUB165-MB4, 6TUB165-MB3 < W-TUB165, Ind-TUB165 < TUB00 < 7TUB165-OC, 7TUB165-MB.

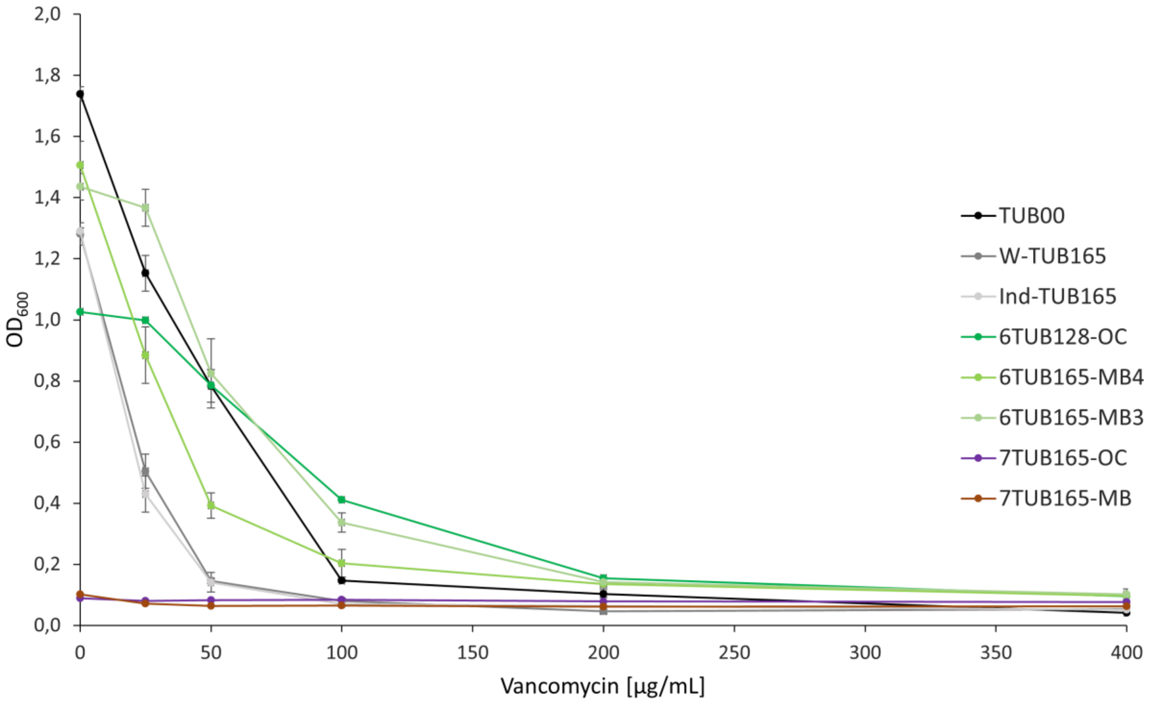

Figure S11. Determination of the MIC of vancomycin. The growth at 24h of the ancestral and all adapted strains treated with a dilution series of vancomycin (0, 25, 50, 100, 200, 400 µg/mL) is shown.

### 12. Expression of 6FTrp- and 7FTrp- substituted variants of EGFP and ECFP in ALE strains

Variants of the green fluorescent protein (GFP) were expressed in the final isolates 6TUB128-OC, 6TUB165-MB4, 6TUB165-MB3 and 7TUB165-OC, 7TUB165-MB. These variants are the C-terminal his-tagged enhanced green fluorescent protein (EGFP-H6) and the N-terminal his-tagged enhanced cyan fluorescent protein (H6-ECFP). The strains were rendered electro competent (by three times washing in 10 % glycerol) and transformed with the respective expression plasmids pQE80L EGFP-H6 or pQE80L H6-ECFP (both endowed with ori ColEI, AmpR).

Inoculated from cryo stocks, precultures of transformed cells were incubated in NMM0-70-0 (6Fi or 7Fi), supplemented with 100 µg/mL ampicillin (Amp) and subsequently 150 mL expression culture (NMM0-70-0 (6Fi or 7Fi), 100 µg/mL Amp) were inoculated to an $OD_{600}$ of 0.05. The cultures were incubated at 30 °C, 180 rpm until they reach exponential growth phase ($OD_{600}$ 0.4 – 0.7). Recombinant protein expression was induced by adding 0.5 mM isopropyl β-D-1-thiogalactopyranoside (IPTG) and performed overnight at 30 °C, 180 rpm.

Cells were harvested by centrifugation (20 min, 10.000 g, 4 °C) and resuspended in B-PER$^{TM}$ (Bacterial Protein Extraction Reagent, Thermo Fisher Scientific) according to manufacturer's instructions. The lysis mixture contained B-PER, lysozyme, DNAse and was incubated 10 - 15 min at room temperature. After centrifugation of cell debris (5 min, 15.000 g, RT), the cell lysate was filtered (Rotilabo PVDF Filter, 0.45 µm pore diameter) and target proteins were purified by using affinity chromatography on Protino Ni-NTA column (1 mL Fast Flow, Machery-Nagel). The purification was carried out using a single-channel peristaltic pump P-1 (VWR) and sodium phosphate-based purification buffer (50 mM Na-P, 300 mM NaCl, two-step imidazole gradient: 20 mM and 40 mM, pH 7.8); followed by elution with 500 mM imidazole. EGFP or else ECFP containing fractions were identified and chromophore maturation was proved by detection of fluorescence by irradiation with a UV lamp at 356 nm. Hereinafter, respective elution fractions were collected, and imidazole was removed by dialysis (Microsep$^{TM}$ Advanced Centrifugal Devices, MCOW 3kDa, Pall Laboratory) against 50 mM Na-P, 100 mM NaCl (pH 7.8). Finally, protein identity and incorporation of 6- and 7-fluorotryptophan was confirmed by LC-ESI-Q-TOF mass spectrometry (Agilent 6530 Accurate-Mass Q-TOF). Protein samples were measured in a concentration of 0.1 mg/mL using the following HPLC parameters: linear gradient from 5 % to 80 % buffer A within 20 min (A: 0.1 % formic acid in MQ-$H_2O$, B: 0.1 % formic acid in acetonitrile), flow rate 0.3 mL/min, injection volume 5 µL. For the MS spectrum a range of 27000 – 29000 amu was selected in the total ion current (TIC) plot and the maximum entropy deconvolution algorithm was applied.

The MS spectra are shown in **Figure S12** and the mass values are shown in Table S1**Table S6**.

6TUB128-OC expressed 2x6FW-ECFP and 1x6FW-EGFP

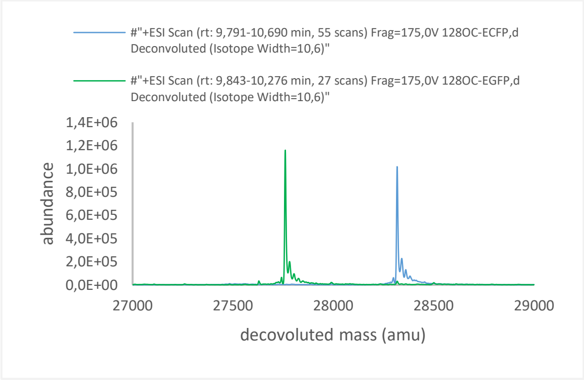

7TUB165-OC expressed 2x7FW-ECFP and 1x7FW-EGFP

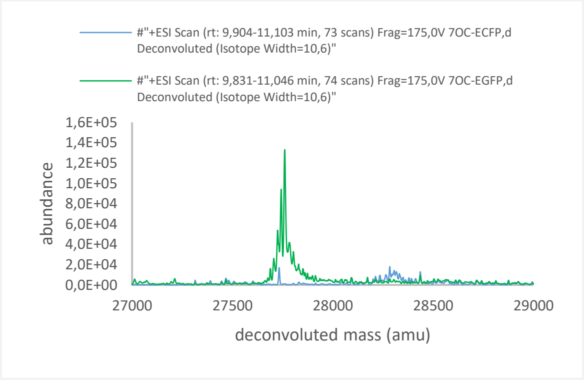

6TUB165-MB4 expressed 2x6FW-ECFP and 1x6FW-EGFP

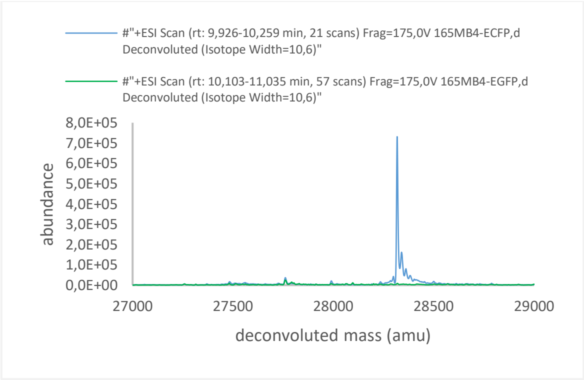

7TUB165-MB expressed 2x7FW-ECFP and 1x7FW-EGFP

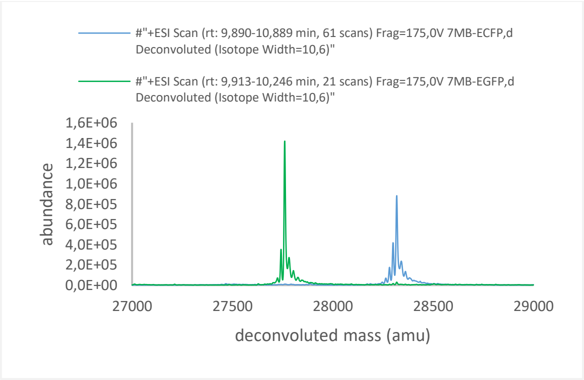

6TUB165-MB3 expressed 2x6FW-ECFP and 1x6FW-EGFP

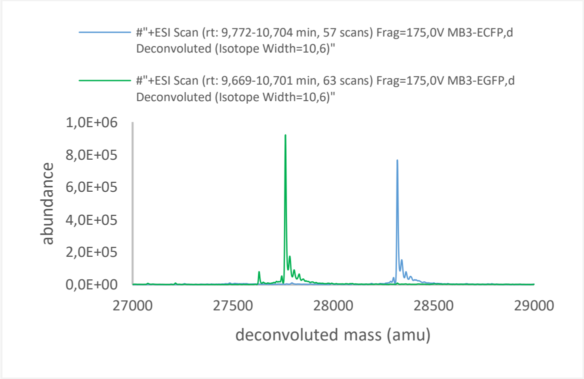

Figure S12. Mass spectrometric analyses of 6FTrp and 7FTrp substituted EGFP and ECFP. Deconvoluted mass spectra of 6FTrp-subsituted ECFP and EGFP expressed in 6Fi-adapted strains are shown left and 7FTrp-substituted ECFP and EGFP expressed in 7Fi-adapted strains are shown right. The ESI scans of EGFP variants are colored in green and of ECFP in blue.

Table S6. MS results of FTrp substituted EGFP and ECFP. Theoretical and observed mass values as well as normalized abundance of EGFP-H6 and H6-ECFP expressed in 6Fi- and 7Fi-adapted strains (after chromophore maturation) are shown.

| | **EGFP (theoretical mass: 27761.56 Da)** | | **ECFP (theoretical mass: 28319.15 Da)** | |
|---|---|---|---|---|
| strain | observed mass [Da] | norm. abundance [%] | observed mass [Da] | norm. abundance [%] |
| 6TUB128-OC | 27761.63 | 81.75 | 28319.36 | 71.72 |
| 6TUB165-MB4 | 27762.29 | 1.73 | 28319.36 | 51.54 |
| 6TUB165-MB3 | 27762.58 | 64.92 | 28320.41 | 54.01 |
| 7TUB165-OC | 27761.71 | 9.38 | nd. | nd. |
| 7TUB165-MB | 27761.44 | 100.00 | 28319.59 | 62.10 |

### References

[1] Datsenko, K. A. and Wanner, B. L. One-step inactivation of chromosomal genes in Escherichia coli K-12 using PCR products. *Proc. Natl. Acad. Sci. U. S. A.* **97,** 6640–6645 (2000).

[2] Crawford, I. P., Nichols, B. P. and Yanofsky, C. Nucleotide sequence of the trpB gene in Escherichia coli and Salmonella typhimurium. *J. Mol. Biol.* **142,** 489–502 (1980).

[3] Wilcox, M. The Enzymatic Synthesis of L-Tryptophan Analogues. *Anal. Biochem.* **440,** 436–440 (1974).

[4] Budisa, N., Steipe, B., Demange, P., Eckerskorn, C., Kellermann, J. and Huber, R. High-level Biosynthetic Substitution of Methionine in Proteins by its Analogs 2-Aminohexanoic Acid, Selenomethionine, Telluromethionine and Ethionine in Escherichia coli. *Eur. J. Biochem.* **230,** 788–796 (1995).

[5] Kublik, A., Deobald, D., Hartwig, S., Schiffmann, C. L., Andrades, A., von Bergen, M., Sawers, R. G. and Adrian, L. Identification of a multi-protein reductive dehalogenase complex in Dehalococcoides mccartyi strain CBDB1 suggests a protein-dependent respiratory electron transport chain obviating quinone involvement. *Environ. Microbiol.* **18,** 3044–3056 (2016).

[6] Seidel, K., Kühnert, J. and Adrian, L. The complexome of Dehalococcoides mccartyi reveals its organohalide respiration-complex is modular. *Front. Microbiol.* **9,** 1130 (2018).
